# Glutamine 666 renders murine ADAM10 an inefficient *S. aureus* α-toxin receptor

**DOI:** 10.1101/2022.05.11.491455

**Authors:** Gisela von Hoven, Martina Meyenburg, Claudia Neukirch, Daniel Siedenschnur, Matthias Husmann

## Abstract

*S. aureus* is one of the most important causes of infectious diseases in hospitalized individuals and outpatients. The majority of clinical isolates secretes large amounts of the small membrane pore-forming α-toxin, alias α-hemolysin, which serves as an important virulence factor of this organism. The identification of *A Disintegrin And Metalloprotease* (ADAM10) as its high affinity receptor held great promise for a better understanding of the processes underlying membrane damage by α-toxin. Twelve years on however, the molecular details of initial toxin binding to target cells remain elusive. Because we noted that several murine cell lines were resilient to α-toxin, we considered the possibility that murine ADAM10 could be less efficient a receptor, as compared to human or bovine orthologues. Accordingly, we sought to identify amino acid residues in ADAM10, which could explain species-dependent functionality as receptor for α-toxin. Our work led to the finding that replacement of a single glutamine residue (Q666) in murine ADAM10 with corresponding glutamic acid (E665) of human or bovine ADAM10 enhances significantly the binding and consequent cytotoxicity of α-toxin. Consistently, a synthetic peptide comprising E665 mitigated α-toxin-dependent hemolysis. In multicellular organisms, E665 is highly conserved, but mice and several other members of the taxon *glires* evolved glutamine at the corresponding position. The residue is located in a short membrane proximal, extracellular region of ADAM10. Taken together, available structural information, *in silico* docking, and functional data suggests that α-toxin monomers could bind to cellular membranes via this so-called stalk region of ADAM10 *and* phosphocholine.

## Introduction

*S. aureus* is among the most common bacterial agents responsible for diseases in humans, ranging from minor infections of the skin to osteomyelitis and life-threatening conditions like pneumonia or septicemia. Consequently, *S. aureus* poses a permanent and huge burden to both patients and health care systems. Thus, in-patients with *S. aureus* infection had, on average, three times the length of hospital stay, three times the total charges, and five times the risk of in-hospital death than inpatients without this infection in the United States (Noskin, Rubin et al. (2005). *S. aureus* α-toxin is an eminent virulence factor of *S. aureus* and archetype of the small β-barrel pore forming toxins (Berube and Bubeck Wardenburg 2013, von Hoven, Qin et al. 2019).The ∼33kD monomer can assemble into a stable heptameric complex forming a water-filled transmembrane channel (Song, Hobaugh et al. 1996). Membrane perforation may lead to hemolysis and cell death in susceptible cells. Despite hundreds of publications on the process of pore formation or its biological consequences, even some of the most fundamental aspects of α-toxin keep puzzling scientists in the field (Berube and Bubeck Wardenburg 2013, von Hoven, Qin et al. 2019). One of the major open questions is how exactly α-toxin interacts with target cells. Early work suggested that high affinity binding sites exist on rabbit red blood cells (RBBC) (Cassidy and Harshman 1973, Hildebrand, Pohl et al. 1991). High-resolution structures of the heptameric α-toxin pore bound to glycerophosphocholines revealed a role of amino acid residues W179 and R200 for the interaction of α-toxin and phosphocholine head groups (Galdiero and Gouaux 2004). Valeva et al. discussed that clustered phosphocholine head groups could serve as high affinity receptor for α-toxin (Valeva, Hellmann et al. 2006). Four years later, Wilke and Bubeck-Wardenburg proposed that *A disintegrin and metalloprotease* 10 (ADAM10) functions as “the probable high affinity α-toxin receptor” (Wilke and Bubeck Wardenburg 2010). By using non-lytic α-toxin mutant H35L fused to glutathion-S-Transferase, the authors had isolated ADAM10 from rabbit red cell ghost preparations. Further, they provided evidence that ADAM10 critically contributes to α-toxin-mediated cellular injury. The suggestion that binding of α-toxin directly activates the catalytic activity of ADAM10 (Inoshima, Inoshima et al. 2011), was not confirmed by others (Ezekwe, Weng et al. 2016, von Hoven, Rivas et al. 2016). However, various groups have provided additional evidence for a role of ADAM10 (Popov, Marceau et al. 2015, von Hoven, Rivas et al. 2016), and/or sphingomyelin (Schwiering, Brack et al. 2013, Virreira Winter, Zychlinsky et al. 2016, Ziesemer, Möller et al. 2019) for the interaction of α-toxin with cellular membranes. Yet, composition and structure of the presumed complex of α-toxin and receptor(s) on target cell membranes are unknown. Recently, the crystal structure of a large portion of the ADAM10 ectodomain has been solved (Seegar, Killingsworth et al. 2017), but so far, this information did not translate into a model of α-toxin/ADAM10 interaction. Using truncation analysis and mutagenesis experiments we found that the ADAM10 prodomain (PD), which acts as a chaperone (Anders, Gilbert et al. 2001), and the disintegrin-domain, are required to efficiently mediate α-toxin-dependent cytotoxicity, whereas an intact catalytic site was dispensable (von Hoven, Rivas et al. 2016).

In cell culture based assays, several murine cell types were rather insensitive to α-toxin (Walev, Martin et al. 1993, Walev, Palmer et al. 1994, von Hoven, Neukirch et al. 2015). Thus, in murine keratinocytes (PDV cells) or murine embryonal fibroblasts (MEF), treatment with 300 nM α-toxin did not alter ATP-levels, although it caused a significant drop of ATP in human fibroblasts and keratinocytes (HaCaT) (von Hoven, Neukirch et al. 2015). Several other nucleated human cell types, with the notable exception of granulocytes (Valeva, Walev et al. 1997), are also comparably sensitive to α-toxin (Bhakdi, Muhly et al. 1989, Jonas, Walev et al. 1994, Dragneva, Anuradha et al. 2001). Expression levels, subcellular distribution of receptors, and cell autonomous defense mechanisms may vary between cell lines and significantly impact the susceptibility to α-toxin (Maurer, Reyes-Robles et al. 2015, von Hoven, Neukirch et al. 2015). In addition, orthologues of ADAM10 or other α-toxin-binding proteins could be of differential efficiency as receptors, but a systematic investigation of this aspect is lacking. Reasoning that species-dependent differences could help to identify regions of ADAM10 critical for its presumed role as a receptor for α-toxin, we decided to address this issue.

## Results

### Is murine ADAM10 an efficient receptor for *S. aureus* α-toxin?

Initially, we compared the susceptibility of the human keratinocyte cell line HaCaT and murine keratinocyte cell line PDV to α-toxin. We measured by flow cytometry the binding of annexin-V, a marker for phosphatidyl-serine (PS)-exposure in the outer leaflet of the plasma membrane, and staining for propidium iodide (PI) as a marker for membrane permeabilization. This experiment revealed massive PS exposure, and PI influx in a subpopulation of PS-positive cells. In stark contrast, murine keratinocytes (PDV) remained virtually unaffected by α-toxin (Figure 1A), in line with previous observations (Walev, Martin et al. 1993, Walev, Palmer et al. 1994, von Hoven, Neukirch et al. 2015). Since many parameters could co-determine differential susceptibility of these two cell lines for α-toxin, it is not trivial to clarify the contribution of differences between ADAM10 orthologues. Therefore, we resorted to HAP1ADAM10KO cells (HAP1 cells with an ADAM10 knock out), (von Hoven, Rivas et al. 2016), as a homogenous system to study the effect of transiently transfected ADAM10 constructs. Previously, we had found that bovine ADAM10 (bADAM10) and human ADAM10 (hADAM10) were of similar efficiency in sensitizing HAP1ADAM10KO cells for α-toxin (von Hoven, Rivas et al. 2016). Here, for comparison of murine and bovine ADAM10, we transiently transfected expression plasmids pcDNA3-mADAM10 and pcDNA3-bADAM10-HA into HAP1ADAM10KO cells, exposed them to α–toxin and measured cellular ATP-levels after 2 hours. We observed a significant drop of ATP in cells receiving the bovine ADAM10 expression construct, but only statistically insignificant reduction in samples transfected with the murine ADAM10 expression plasmid (Figure 1B). To confirm expression of ADAM10 in transfected cells we used Western-blots with an antibody raised against a C-terminal peptide (732-748), of human (or bovine) ADAM10. Reportedly, the antibody can bind to murine ADAM10 as well. However, HAP1ADAM10KO cells transfected with murine ADAM10 yielded weak bands of dubious specificity (Figure 1C), raising the question whether expression-and/or detection efficiency was/were low. Because in the murine sequence of the C-terminal peptide one amino acid (residue 734) differs from the bovine sequence (supplementary Figure 1), we could not exclude that detection of murine ADAM10 with this antibody was less sensitive than detection of bovine ADAM10.

**Fig.1.**
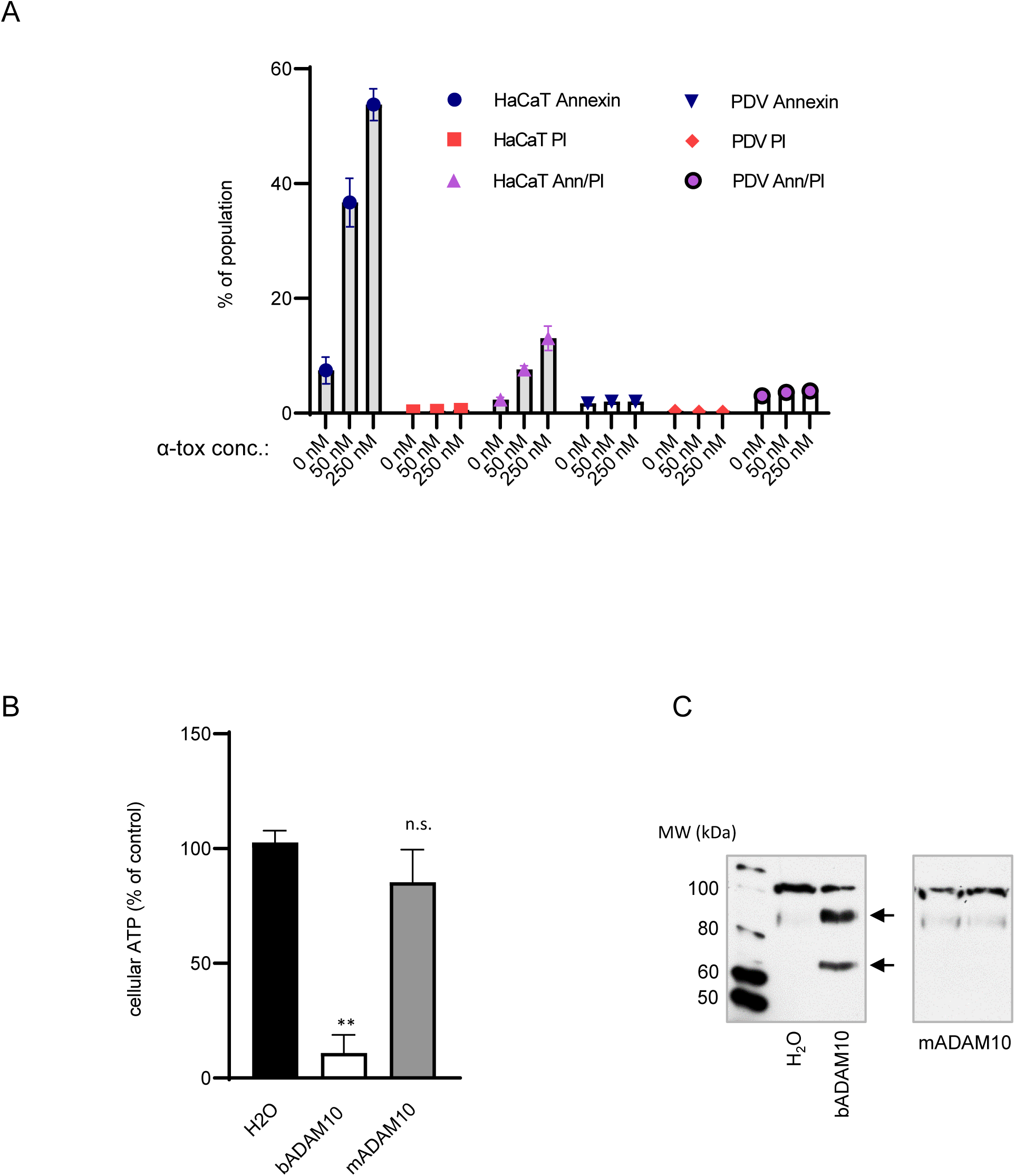
Indication that mADAM10 could be a comparably inefficient α-toxin receptor. (A) Human keratinocytes (HaCaT), or mouse keratinocytes (PDV) were left untreated or incubated with *S. aureus* α-toxin (50 nM or 250 nM) for 2h, washed, and analyzed by FACS for exposure of phosphatidylserine at the outer leaflet of the plasma membrane (staining with annexinV-AlexaFluo647, Invitrogen, A23204), and for membrane damage, using PI. Bars indicate percent positive cells (Annexin V, PI, both). Data are mean values from four independent experiments, error bars indicate ± SEM. (B) HAP1ADAM10KO cells were mock transfected (H_2_O), or transiently transfected with plasmids encoding for bovine or murine wild type ADAM10. The next day quadruplicate samples were treated, or not, with 500 ng/ml α-toxin (16.6 nM) for 2h, before measuring cellular ATP. Shown are mean values of three independent experiments ± SEM; ** indicates significance between expression-plasmid transfections and mock transfection (H_2_O), with an adjusted *P* value of 0.0079 in a one-way ANOVA and Dunnett’s multiple comparisons; no significance (n.s.) was found for the comparison of mADAM10 vs. mock transfection. (C) Western-blot for detection of ADAM10 (antibody A124695) in whole cell lysates from cells, transfected as in (B); arrows on the right side indicate processed ADAM10 (upper), and mature ADAM10, respectively; note strong bands in samples with bADAM10 only.

### Apparent inefficiency of mADAM10 as α-toxin receptor does not reside in the PD

Hence, for reliable comparison of function and expression of these orthologues, we used plasmids encoding murine or bovine ADAM10-derivatives, carrying C-terminal hemagglutinin (HA)-tags. For expression of ADAM10, we co-transfected HAP1ADAM10KO cells with separate plasmids encoding for either PD or ΔPD. This allowed us to perform a swapping experiment aimed at localizing potential species-dependent differential functionality to either the PD or the ΔPD. PD and ΔPD, when co-expressed from separate plasmids do function like intact ADAM10 (Anders, Gilbert et al. 2001, von Hoven, Rivas et al. 2016). Initially, we favored the idea that the murine PD might account for the presumed inefficiency of murine ADAM10 as a receptor for α-toxin, because first, the longest contiguous stretch of mismatches between ADAM10 from *mus musculus* and *bos taurus* (and *homo sapiens*) is located in the C-terminal part of the PD (residue 193-198), (supplementary Figure 1). Second, we had previously found that expression of the PD of bADAM10 is important for α-toxin-dependent cytotoxicity (von Hoven, Rivas et al. 2016). Third, two rare mutations in the ADAM10 PD, (Q170H and R181G), which impair the chaperone function of the PD (Suh, Choi et al. 2013), and co-segregate with Late Onset Alzheimer’s Disease (LOAD), (Kim, Suh et al. 2009), appeared to enhance α-toxin-dependent cytotoxicity in preliminary experiments (supplementary Figure 2). Therefore, we co-expressed murine or bovine PD with either bovine or murine ΔPD as illustrated in Figure 2A, and confirmed expression by Western-blot (Figure 2B). Next, we treated co-transfected cells with α-toxin and measured cellular ATP. This experiment revealed that murine ΔPD, whether co-transfected with murine or bovine PD, failed to efficiently confer α-toxin-dependent loss of ATP (Figure 2C). In contrast, the bovine ΔPD mediated α-toxin-dependent loss of ATP, in conjunction with either bovine or murine PD. This led to three conclusions: first, murine ADAM10 is probably not an efficient mediator of α-toxin-dependent cytotoxicity, second, the superior function of bADAM10 in this regard resides in the ΔPD-part of the protein, and third, murine or bovine PDs are exchangeable in this context.

**Fig.2.**
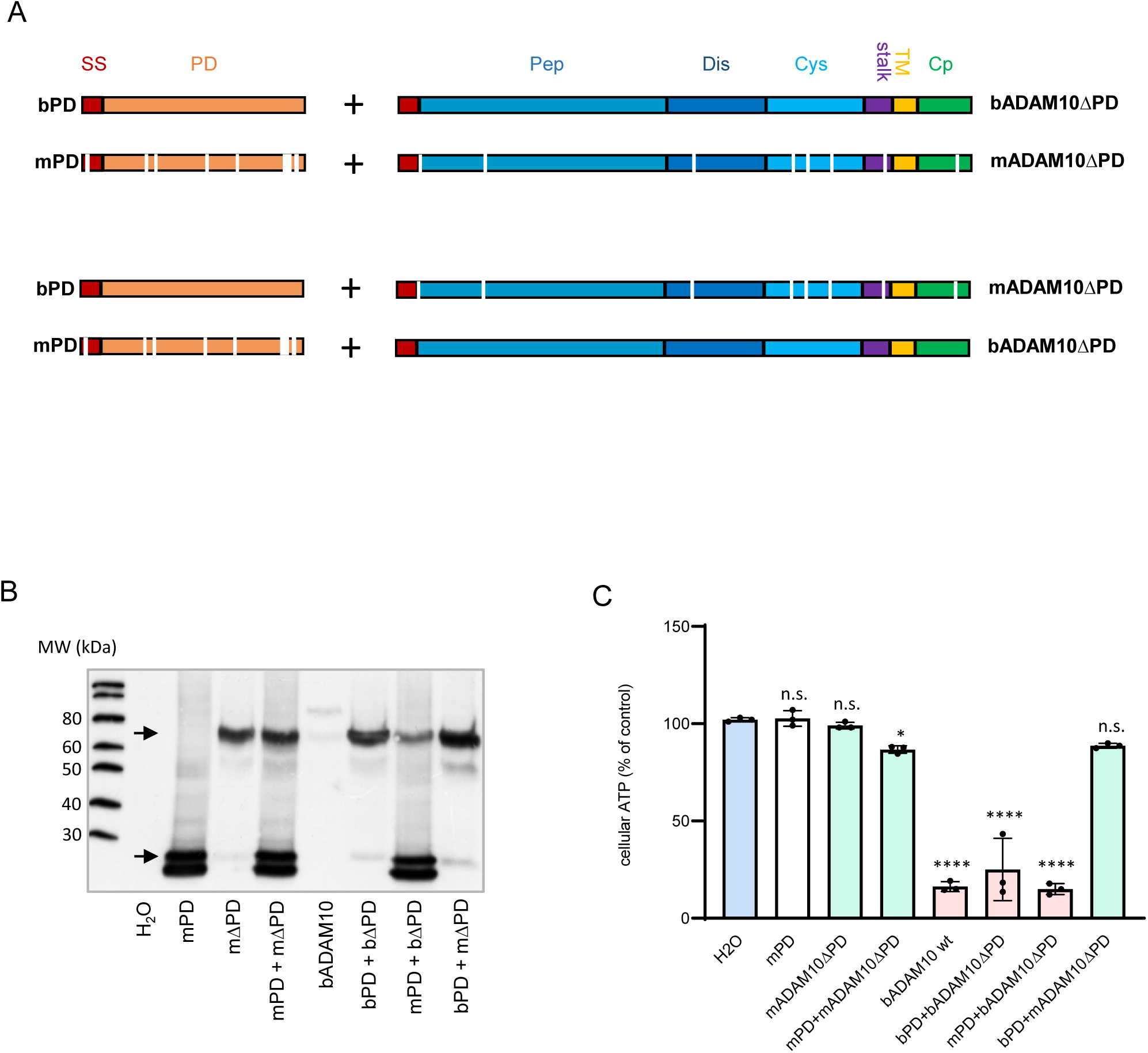
The α-toxin-resilient phenotype of murine ADAM10 resides in the ΔPD. (A) Schematic of co-expression experiments with murine or bovine PD, in combination with either cognate ΔPD (upper part of the figure), or with orthologous ΔPD (swapping experiment, lower part of the figure). The signal sequence (SS) is labeled in red, the prodomain (PD) in orange, the peptidase-(Pep), disintegrin-(Dis) and cystein-rich-domain in different hues of blue, stalk-region (*stalk*) in purple, transmembrane domain (TM) in yellow and the cytoplasmic domain (Cp) is labeled in green. All constructs (PD and ΔPD) carry a HA-tag at their C-termini, omitted from the schematic for clarity. White markings indicate positions where amino acids differ in the mouse sequence as compared to both bovine and human sequences. (B) Western blot analysis of lysates from HAP1ADAM10KO cells mock-transfected or (co-)transfected as indicated in the figure, using antibody against the HA-tag. Arrows point to ΔPD (upper), and PD (lower). (C) ATP-levels in lysates from HAP1ADAM10KO cells mock-transfected, or (co)-transfected as indicated in the figure, and treated with 500 ng/ml α-toxin for 2h. Data represent ATP-levels of α-toxin treated samples vs. untreated for each transfection mix. Shown are mean values of three independent experiments ± SEM. **** indicates adjusted *P* Values < 0.0001, and * an exact *P*-value of 0.037 for comparisons of transfected vs. H_2_O control (one-way ANOVA with Dunnett’s multiple comparisons test).

### Residue E665 is important for the function of bADAM10 as an efficient α-toxin receptor

Next, we sought to identify regions or residues in the ΔPD responsible for the different efficiency of murine vs. bovine ADAM10 as mediator of α-toxin-dependent cytotoxicity. There are eight amino acid residues in the ΔPD, which are conserved in human and bovine ADAM10, but different in the mouse. These residues are scattered over all domains (i.e. the Met-, Dis-, Cys-, Tra-and Cyt-domain, whereby acronyms stand for Metalloprotease-, Disintegrin-, Cystein-rich-, Transmembrane-and Cytosolic domain), (Figure 3A). Any combination of (any subset of) these eight amino acid residues could be responsible for the comparably low susceptibility of cells expressing mADAM10 to α-toxin. However, we considered it possible that few residues, or even a single one, might be responsible for the functional difference between mADAM10 and bADAM10. In an attempt to narrow in on relevant residues, we studied the function of two *in vitro* synthesized bovine ΔPD derivatives, each with four different replacements out of the eight “mouse-specific” residues indicated in Figure 3B. When co-transfected with PD, a construct comprising mutations at position N°2, N°3, N°4 and N°6 mediated toxicity similar to bovine wild type ΔPD, co-transfected with PD (Figure 3C). Fortuitously, the second construct, comprising mutations at positions N°1, N°5, N°7 and N°8, when co-transfected with PD, failed to render HAP1ADAM10KO cells susceptible to α-toxin (Figure 3C). Consequently, we generated three bovine ΔPD constructs each with single residues (at positions N°5, N°7 or N°8, as shown in Fig. 3B) mutated to the mouse sequence. Strikingly, mutation at position N°7, (E665Q), but not at position N°8, (Q734P), was sufficient to deprive the bovine ΔPD construct of its ability to confer α-toxin-dependent cytotoxicity (Figure 3C). Mutation N°5 (D591N) also failed to affect ADAM10’s function as a mediator of α-toxin-dependent toxicity (data not shown). E665 in bADAM10 corresponds to Q666 in mADAM10; the shift in numbering is due to an additional amino acid (serine) in the sequence of mADAM10, following residue 203 (supplementary Figure 1).

**Fig.3.**
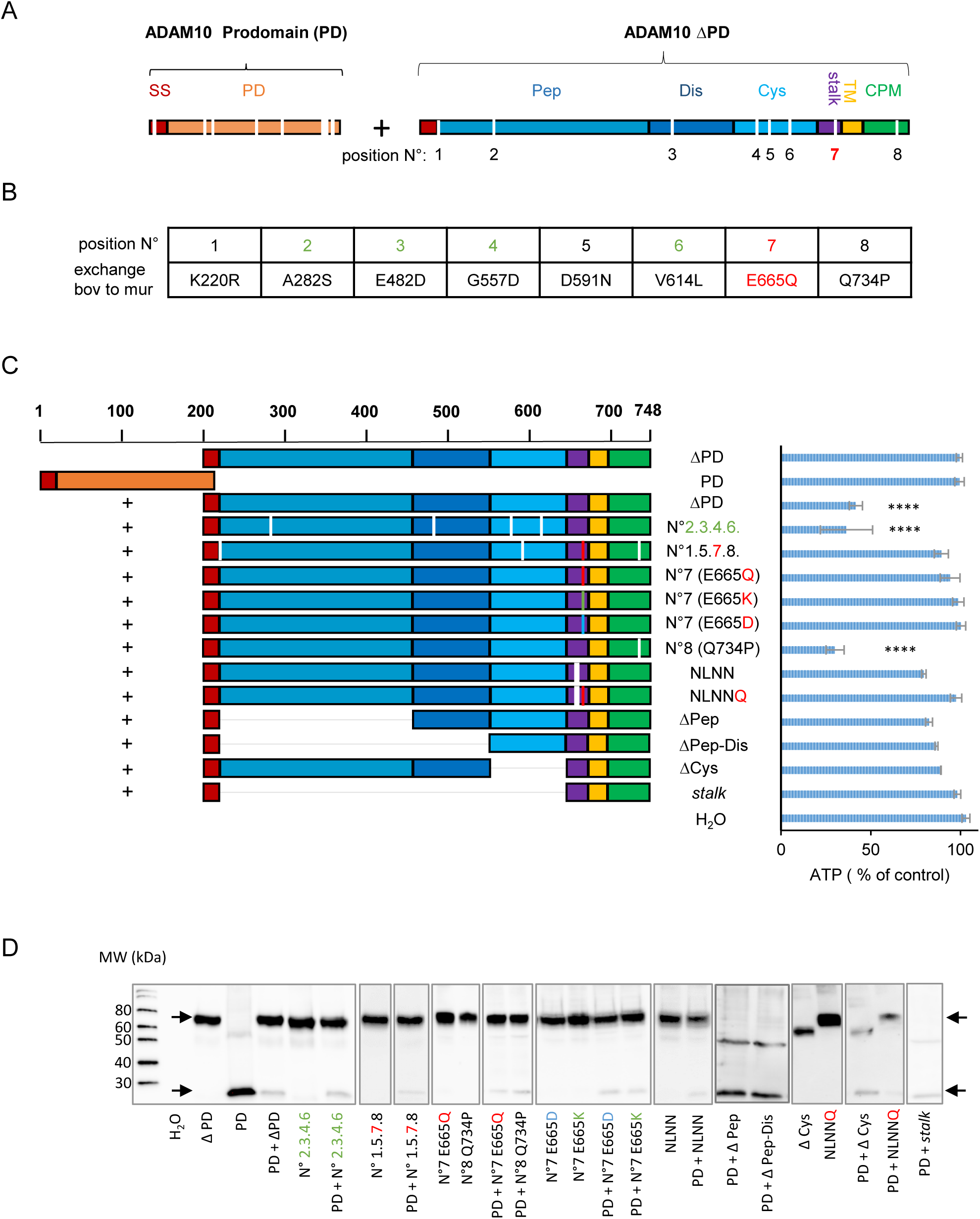
E665 of bADAM10 is important for cellular susceptibility to α-toxin. (A) Position N° 1 through 8 indicated below ADAM10ΔPD in this schematic highlight the eight amino acid residues in the mouse sequence, which deviate from both bADAM10 and hADAM10. (B) The table specifies the amino acid exchanges at positions numbered 1-8 in (A). Green numbers indicate those positions, changed in one of two compound mutants, black numbers, and the red “7” (corresponding to E665Q), indicate positions changed in the second compound mutant. (C) Left: Schematic showing bADAM10 truncation constructs, compound mutations or single residue mutations tested for their ability to confer to HAP1ADAM10KO cells susceptibility to α-toxin, when co-transfected with bovine PD (indicated by “+” on the left side). Samples receiving PD or ΔPD alone (two uppermost constructs) served as controls. Scale at the top indicates amino acid residue numbers in bADAM10. Color code of domains is as in (A). Right: Bars indicate ATP levels in HAP1ADAM10KO cells, which were (co)-transfected as indicated in the figure and treated with 500 ng/ml α-toxin for 2h relative to samples receiving no toxin. Data are from n ≥ 3 independent experiments; mean values ± SEM. Mutations at position N°7 (corresponding to E665 in bADAM10) are highlighted in red; these constructs, when co-transfected with PD, failed to confer to HAP1ADAM10KO cells sensitivity to α-toxin; the difference to H_2_O was not significant in a one way ANOVA with Tukey’s multiple comparisons. Similarly, no significant differences were found for NLNN, NLNNQ, ΔPep, ΔPep-Dis, ΔCys and *stalk*. In contrast, the compound mutation with changes at positions N° 2.3.4.6 (highlighted in green) plus PD, or mutant N°8 plus PD were fully active, like wild type ΔPD plus PD. **** indicates *P* < 0.0001 for comparisons with H_2_O control. (D) Western-blot using antibodies directed against the HA-Tag, for verification of expression of ADAM10-constructs indicated in the figure; arrows indicate ΔPD(-variants) (upper arrow), and PD (lower).

At physiological pH, Q is usually electro-neutral, whereas E has a negative charge. Therefore, it was of interest to see whether replacement of E665 with other charged amino acids, affects the function of ADAM10 as a mediator of α-toxin-dependent cytotoxicity. Mutant bADAM10 E665<u>K</u>, where lysine, which has a positive charge, replaces glutamic acid proved to be inefficient as mediator of α-toxin action (Figure 3C). Substitution of E665 with D, thus retaining the negative charge, too failed to confer sensitivity to α-toxin (Figure 3C). Therefore, size and orientation of the amino acid side chain at position 665, possibly in addition to its negative charge, could be important for optimal function of bADAM10 as a receptor for α-toxin. We verified equal expression of all constructs by Western-blot using antibodies against the HA-tag at the C-terminus (Figure 3D). Further, we compared expression of mutant ΔPD E665Q (N°7), with mutant Q734P (N°8) by FACS analysis, and found both of them equally expressed on the cell surface (Figure 4).

**Fig.4.**
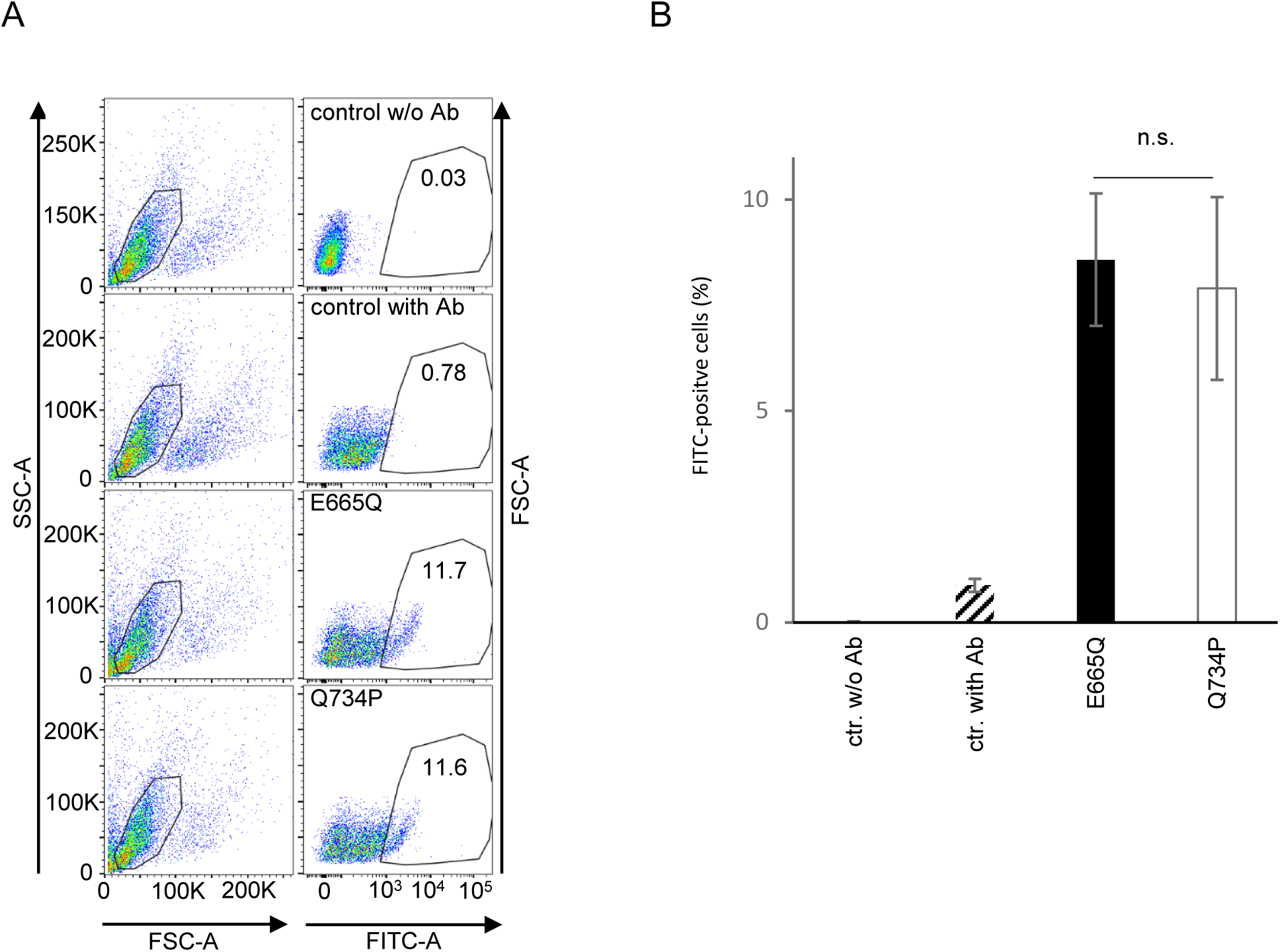
E665Q and Q734P are equally well expressed on the cell surface. (A) Plot of FACS-analysis of HAP1ADAM10KO cells transfected with bADAM10E665Q or bADAM10Q734P and subsequently stained for surface-expression of ADAM10 as detailed in the Methods section. Plots on the left hand show forward scatter *vs*. side scatter, plots on the right hand fluorescence intensity (FITC-A, log scale) vs. number of events. The two uppermost dot plots are controls w/o antibody, or with antibody, but no transfection of ADAM10. The two lower plots on the right hand document similar surface expression of bADAM10E665Q and bADAM10Q734P. (B) Summary of data from three independent experiments as in (A). Mean values ± SEM. The difference between the two constructs was not significant in a two-tailed *Student’s* t-test.

### The stalk region is required for the function of ADAM10 as mediator of α-toxin-dependent cytotoxicity

Residue E665 is located in a membrane-proximal region of ADAM10, which has been termed “stalk-region.” Bleibaum et al. implicated this part of ADAM10 in cell stress-dependent activation of the enzyme (Bleibaum, Sommer et al. 2019). They proposed that activation leads to relocation of the stalk region through interaction with phosphatidylserine residues exposed at the cell surface after calcium ion influx into cells. The proposed model envisages binding of positively charged (K and R) amino acid residues of the stalk region to PS, because replacing RLKK, corresponding to residues 656-659 in bADAM10, with NLNN greatly diminished binding to the phospholipid (Bleibaum, Sommer et al. 2019). As shown here, the same mutations of the stalk region rendered ADAM10 less active a mediator of α-toxin toxicity, as measured by ATP-loss; the effect was stronger with additional mutation E665Q (the construct is designated NLNNQ – with Q in red color - in Figure 3C). The collective results indicate that the stalk region is important for ADAM10’s function as mediator of α-toxin-dependent toxicity. However, a construct with a deletion of the ectodomain except for the stalk-region, was not sufficient to confer toxicity of α-toxin in HAP1ADAM10KO cells (Figure 3C). This is in keeping with our earlier observation that deletion of the disintegrin-domain significantly reduces the ability of ADAM10 to mediate α-toxin-dependent toxicity (von Hoven, Rivas et al. 2016). Although our earlier results had uncovered that an intact catalytic site is not required, it remained possible that the peptidase domain is important for interaction with α-toxin. Accordingly, we generated bADAM10 truncation products lacking the protease domain (ΔPep), or both the protease and the disintegrin domain (ΔPep-Dis); both constructs barely conferred toxicity (Figure 3C). Finally, we generated a construct without Cys-domain (ΔCys), and this too failed to enhance susceptibility of cells to α-toxin (Figure 3C).

### A peptide, covering residue E665 of bADAM10 reduces α-toxin-dependent hemolysis

An urgent question arising from the above results was whether E665 might be directly involved in binding of α-toxin to ADAM10. If this was the case, peptides covering this residue might compete with ADAM10 for binding to α-toxin, and inhibit pore formation. We chose to measure hemolysis as functional read out. The assay is simple and rapid, and it requires only small volumes of reagents, thus minimizing amounts of peptide required. RRBC readily lyse in the presence of nanomolar concentrations of α-toxin. Suitable synthetic peptides can inhibit protein/protein interactions (Bruzzoni-Giovanelli, Alezra et al. 2018). Here we used a custom-made, synthetic peptide covering the wild type sequence SP<u>E</u>LYENIA (E665 underlined) from the stalk region of bADAM10 (Figure 5A). This peptide, but not a control peptide with a replacement of E665 by lysine (K) (SP<u>K</u>LYENIA), reduced lysis by up to 56% (Figure 5B), compatible with the idea that this region may be involved in interactions with α-toxin. The published high-resolution structure of the ADAM10 ectodomain does not cover the stalk region implicated here in mediating cytotoxicity of α-toxin. So we extended the primary sequence of the part of hADAM10, which has been used for structural analysis of the ectodomain (Seegar, Killingsworth et al. 2017) by the sequence of the stalk region and submitted this sequence for structure prediction by the I-TASSER server (Yang, Yan et al. 2015). Not surprisingly, the predicted structures of the stalk region differed substantially even among the five top scoring models, whereas the bulk of the ectodomain matched the published structure. In the present context, we considered the model depicted in Figure 5 C of particular interest, because here, E665 seemed to be potentially accessible for α-toxin.

**Fig.5.**
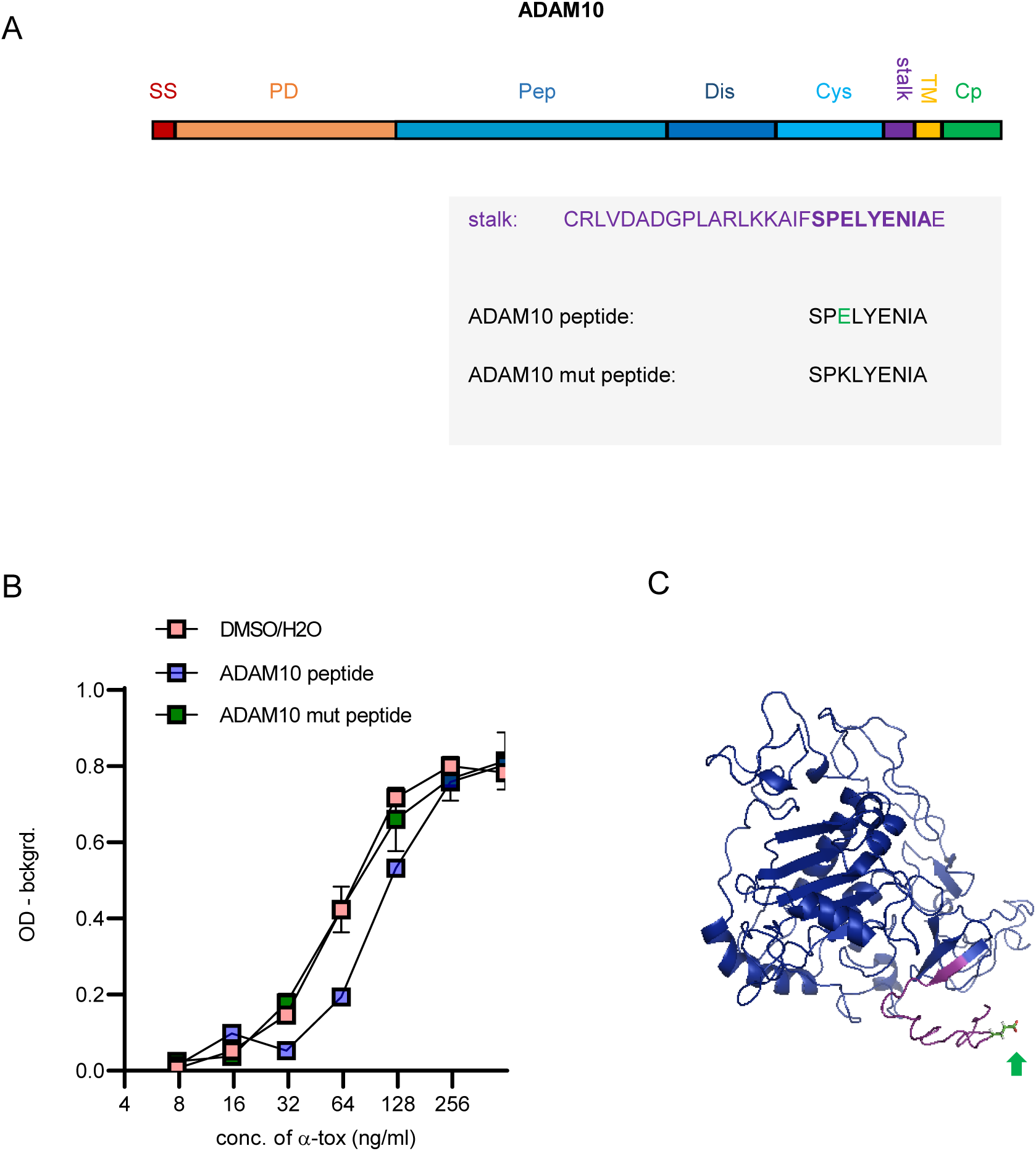
A peptide from the stalk region of bADAM10 reduces α-toxin-dependent hemolysis. (A) Grey box below the schematic of ADAM10 domain organization displays the sequence of the *stalk (*purple) region and sequences of wild type (wt) and mutant ADAM10-derived peptides below. “E” labeled in green in the wt peptide sequence corresponds to residue E665 of hADAM10 or bADAM10. In mutant peptide, glutamic acid (E) was changed to lysine (K). (B) hemolysis of RRBC, incubated for 30 min at 37°C with indicated doses of *S. aureus* α-toxin in the presence of solvent or peptides (10 μM). Data are background-substracted OD. Mean values ± SEM from n=6 independent experiments. At α-toxin concentrations of 125 nM, 62.5 nM and 31.25 nM, hemolysis was significantly lower in the presence of wild type peptide as compared to samples receiving mutant peptide; adjusted *P*-values: 0.011, 0,015 and 0.03, respectively (two-way ANOVA and Tukey’s multiple comparison test). (C) Hypothetical model of ADAM10-ectodomain (hADAM10) including the stalk region (magenta); green arrow points to the side chain of E665, highlighted in green. The model was created using the I-TASSER server (Yang, Yan et al. 2015), see Methods section. A high-resolution structure of a large portion of the ectodomain has been published (Seegar, Killingsworth et al. 2017), and the corresponding part of the model of the present figure essentially matches that structure; the structure of the stalk region is uncertain.

### A hypothetical model of α-toxin/ADAM10 interaction

Because E665 of ADAM10 was required for efficient α-toxin-dependent cytotoxicity and because a peptide covering this residue reduced α-toxin-dependent hemolysis, it appeared possible that this residue contacts α-toxin. In the absence of structural data of a complex of putative receptor and ligand, we employed *in silico* docking in order to obtain a hypothetical model of α-toxin in complex with ADAM10 (see Methods section). The model shown in Figure 6A could be a reasonable approximation. It would seem to allow for simultaneous interaction of the α-toxin monomer with both E665 of ADAM10 *and* phosphocholine of the plasma membrane, which was shown to interact with residues R200 and W179 in the α-toxin heptamer (Galdiero and Gouaux 2004). By virtue of the negative charge of its side chain, E665 of ADAM10 might establish, a polar contact with S203 of the α-toxin monomer (Figure 6B).

**Fig.6.**
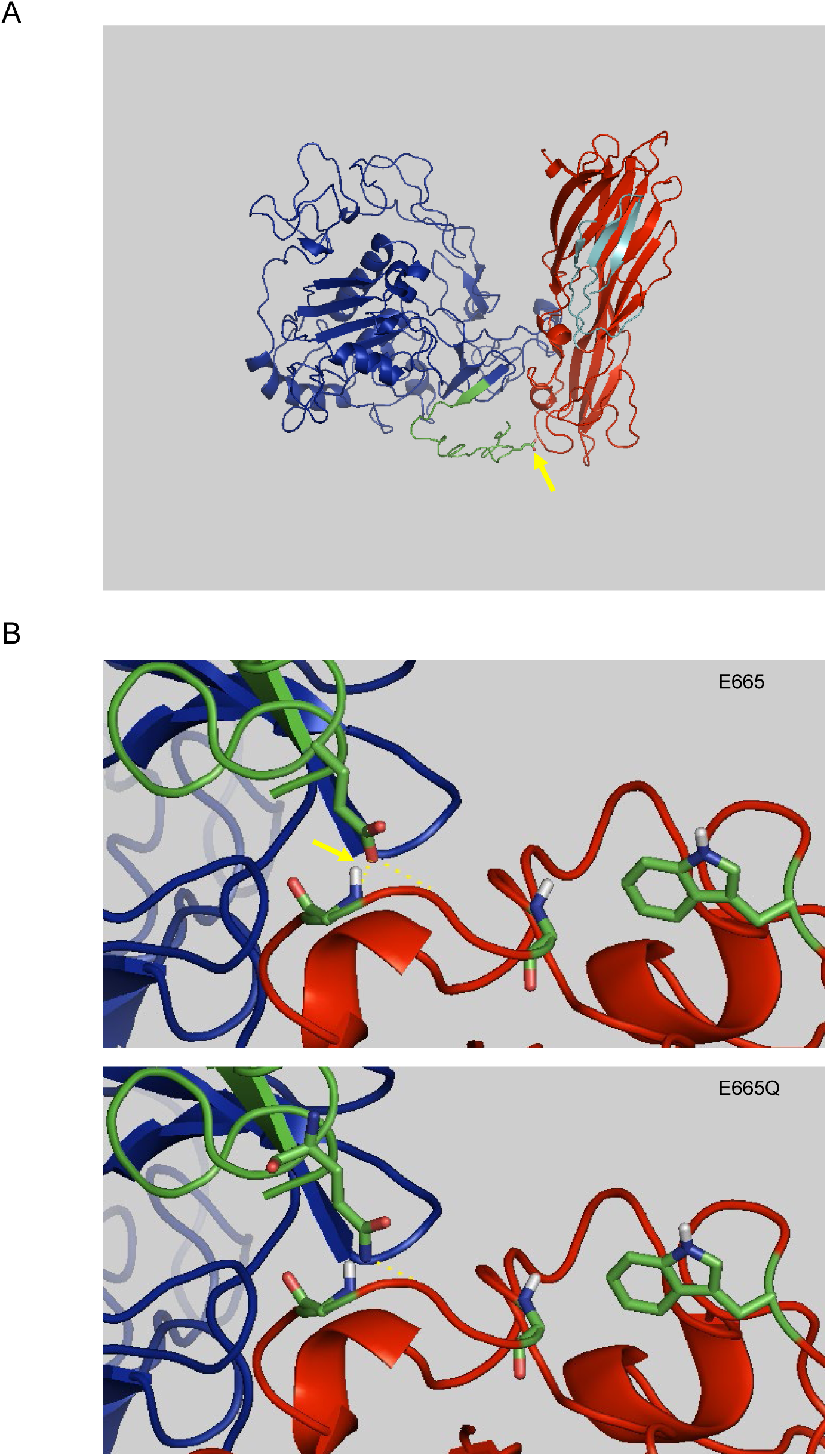
Hypothetical model of ADAM10/α-toxin-interaction. (A) *In silico* docking of α-toxin monomer (red, pre-stem-region in cyan) to the hADAM10 ectodomain (blue) including the stalk region (green). For details see Methods section. The yellow arrow points to the side chain of E665 in the stalk region of ADAM10. (B) Close up view of the base (i.e. the presumed membrane proximal face) of the complex shown in (A). In the upper image of (B) a polar contact is indicated (yellow dotted line highlighted by yellow arrow) between the side chain of E665 in ADAM10 and S203 (main chain indicated) of α-toxin. Side chains of R200 and W179 of α-toxin are on the right hand. Lower image in (B) shows result for *in silico* mutant E665Q.

### Mutation Q666E is sufficient to render mADAM10 a more efficient α-toxin-receptor

Although mutation of a single residue in a short membrane proximal region of bADAM10 significantly diminished the effect of α-toxin, the overall-picture emerging from the data summarized in the text and Figure 3C was that all ADAM10 domains were somehow involved in mediating α-toxin-dependent cytotoxicity. Therefore, we wondered whether mutating Q666 in mADAM10 to E would be *sufficient* to render mADAM10 a more efficient mediator of α-toxin-dependent cytotoxicity. As shown in Figure 7A, this proved to be the case: at 10nM, and more so at 50 nM α-toxin there was a significant difference between mutant and wild type.

**Fig.7.**
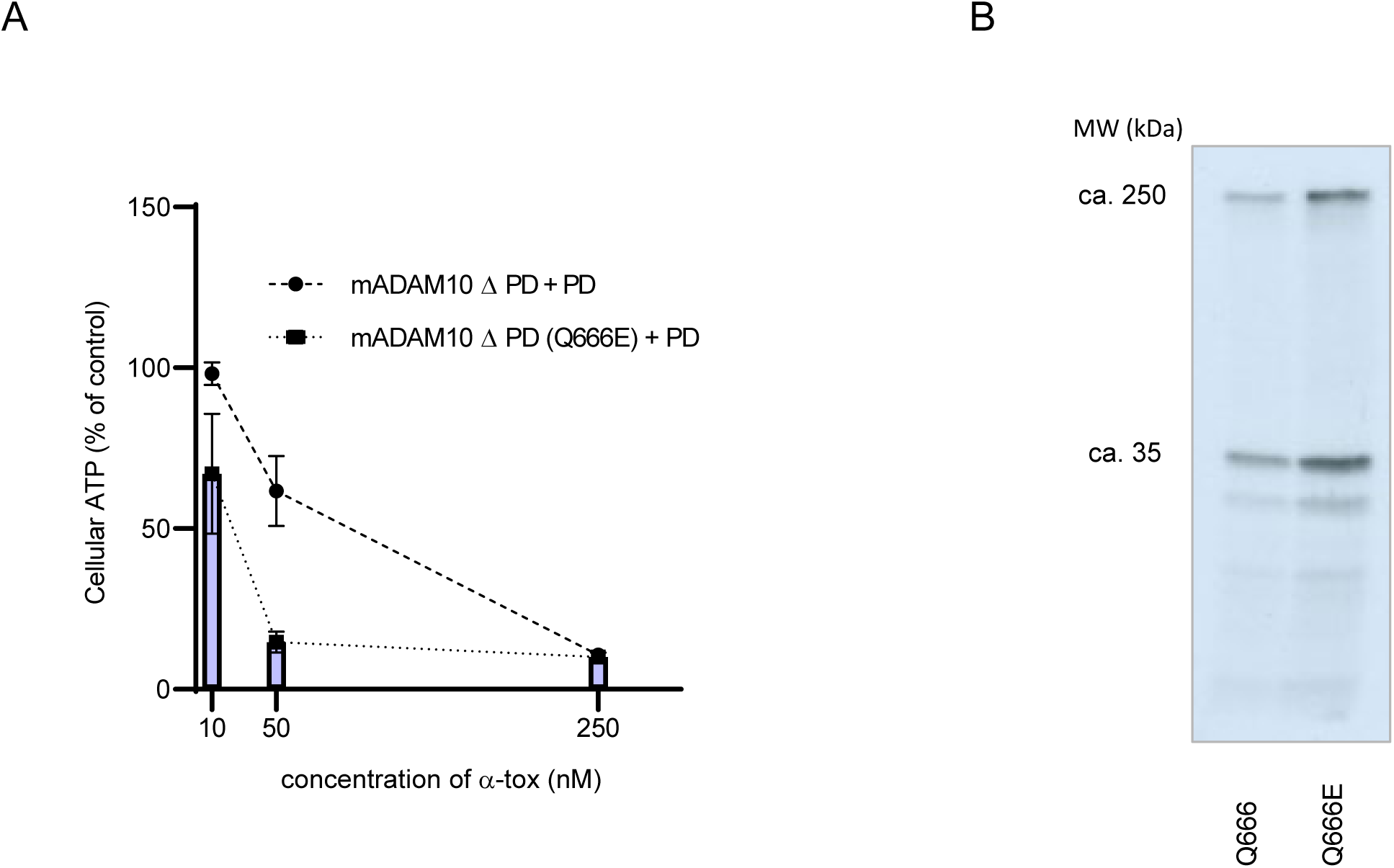
Q666E enhances the efficiency of mADAM10 as α-toxin-receptor. (A) ATP-levels of HAP1ADAM10KO cells transiently (co-)transfected with wild type (Q666) or mutant (Q666E) mADAM10ΔPD plus bPD, and treated, or not, with α-toxin at concentrations indicated in the figure. Data are ATP levels in α-toxin-treated samples relative to samples w/o α-toxin. Mean values of three independent experiments ± SEM. Statistical significance was determined using two-way ANOVA with Sidak’s multiple comparison; adjusted *P*-values for comparing transfections of wild type ADAM10ΔPD vs. ADAM10ΔPD(Q666E) at 10 nM or 50 nM α-toxin indicated a significant difference (0.0038 and 0,0001 respectively), but not at 250 nM α-toxin (*P*=0.9997). (B) The image shows α-toxin monomers (1mers) and SDS-stable oligomers, i.e. putative heptamers (7mers), recovered by immunoprecipitation from lysates of cells co-transfected as in (A) and incubated with internally [^35^S]methionine-radiolabeled α-toxin. Lower molecular weight bands are decay products of labeled α-toxin. Densities of bands corresponding to 7mers and 1mers were 2.0(SEM ± 0.35)-fold and 1.7(SEM ± 0.28)-fold higher for Q666E vs. Q666 (n=3, *P* < 0.05 in two-tailed Student’s t-tests).

Thus far, we had performed functional analyses of truncations and mutations of ADAM10 in order to delineate domains and residues important for ADAM10’s function as a mediator of α-toxin-dependent cytotoxicity. Given that Q666E rendered mADAM10 more efficient in this regard, we sought to investigate the impact of this mutation on binding of toxin and on heptamer-formation. We incubated cells with internally [^35^S]methionine-radiolabeled α-toxin, lysed cells immune-precipitated α-toxin, separated the material by SDS-PAGE and detected [^35^S]methionine-radiolabeled α-toxin by autoradiography. We recovered more toxin from cells transfected with mADAM10-(Q666E), as compared with wild type mADAM10 (Figure 7B). Thus, lower toxin-binding efficiency of mADAM10, as compared with bADAM10, is at least one plausible reason for its lower activity in conferring α-toxin-dependent cytotoxicity.

### A cluster of “Q666-type” ADAM10 among rodents

Alignment and phylogenetic analysis of ADAM10 sequences from metazoan species reveals a high degree of conservation of the residue corresponding to E665 in bADAM10. The rooted tree shown in Figure 8 demonstrates that species as distantly related as frog, fish, birds and mammals, including *H. sapiens*, feature glutamic acid residue at this position; and it is also present in *Saccoglossus kovalevskii*, an acorn worm belonging to the phylum hemichordata (not shown). In spite of the evolutionary conservation of E665, two clusters within the taxon *glires* have evolved a glutamine residue at the corresponding position.

**Fig.8.**
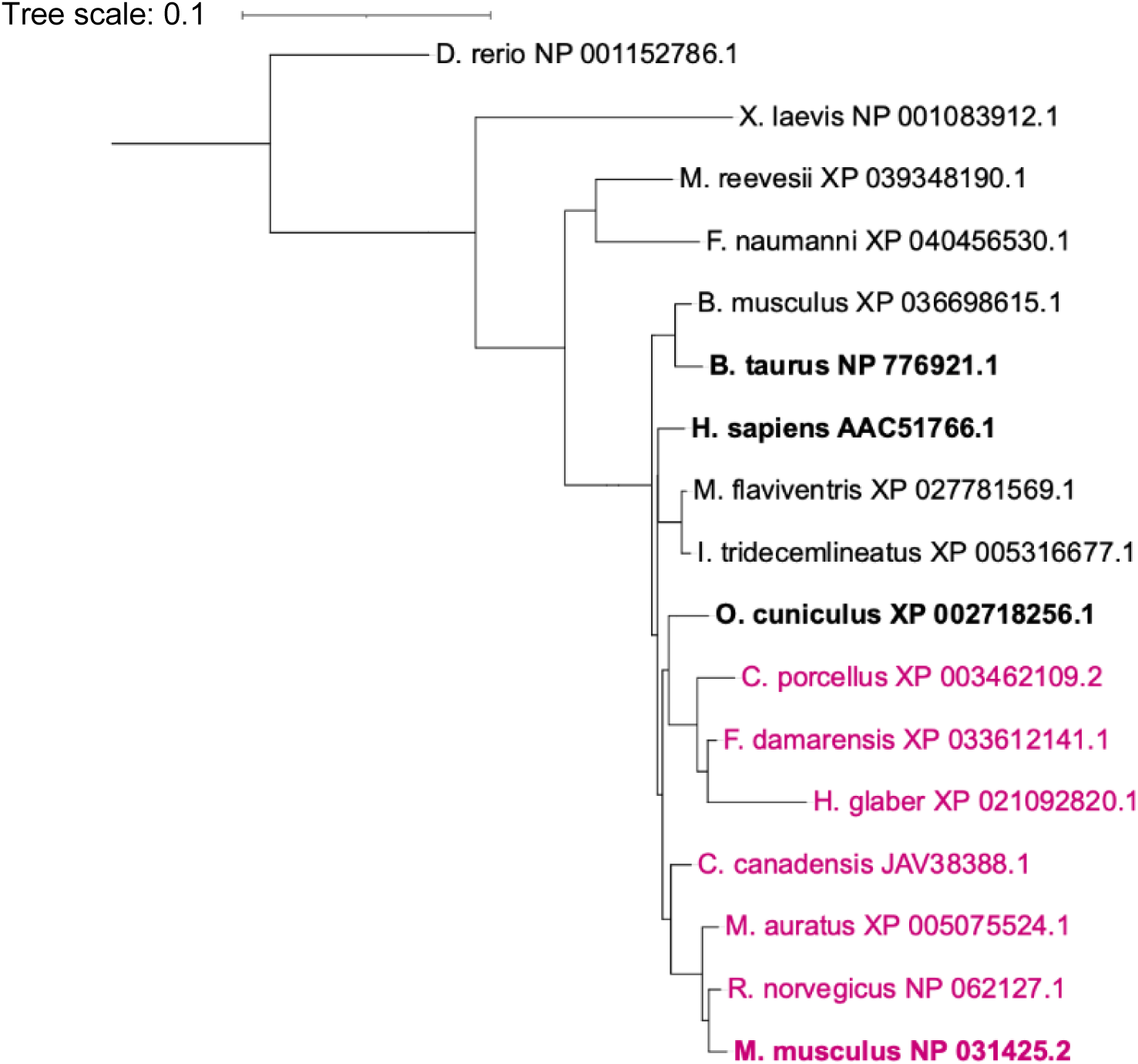
E665 is highly conserved, but some members of the taxon *glires* evolved Q at the corresponding position. Phylogenetic tree based on multiple sequence alignment of ADAM10 protein sequences (supplementary information) of different species; carriers of glutamine Q666 (e.g. mouse, or corresponding position in orthologues) highlighted in magenta.

## Discussion

The principal discovery of the present study is that a single residue (Q666) in the stalk region of murine ADAM10 (mADAM10) can determine species-selective, relative cellular resilience to α-toxin. It offers a straightforward explanation for discrepancies of cell culture experiments with murine cells *vs*. cells from other species. Moreover, it led to a hypothetical model of α-toxin ADAM10 interaction, which accommodates previous and current observations and may explain binding of the α-toxin monomer to cellular membranes with high avidity.

Data from mutation analysis, binding assay and inhibition with a peptide covering E665 of bADAM10 let us hypothesize that this residue contributes to the elusive binding interface of ADAM10 and α-toxin monomer. Satisfyingly, according to the model in Figure 6, the residues changed in mutants N° 2, 3, 4 and 6 (see Figure 3B), which did not affect ADAM10’s ability to mediate α-toxin-dependent cytotoxicity, would probably not interact with α-toxin (supplementary Figure 3). Located in the stalk region of ADAM10, E665 is in proximity to the plasma membrane. Consequently, to interact with E665, residues in α-toxin would have to be similar close to the lipid bilayer. From *in silico* docking using the HDOCK server without restraints, S203 of α-toxin emerged as a potential interaction partner for E665, because this residue could be in sufficient proximity to the negatively charged side chain of ADAM10’s E665 to establish a salt bridge.

Previous work by others showed that R200 in the rim domain of the α-toxin heptamer interacts via a water molecule with the negative charge of phosphocholine head groups, whereas T179 contacts the positive charge of the quaternary ammonium group (Galdiero and Gouaux 2004). Presumably, these interactions will also occur on cellular membranes, and it is possible that phosphocholine can also interact with R200 and T179 of monomers. This prompts the question whether α-toxin monomer bound to phosphocholine could simultaneously interact with its putative high affinity receptor ADAM10. The model shown in Fig. 6 seems to be compatible with such dual binding. As suggested previously, the first α-toxin monomer which binds to a given ADAM10-molecule could provide a binding interface for the next monomer and so forth, whereby all α-toxin monomers would also contact phosphocholine head groups (von Hoven, Qin et al. 2019).

Our preliminary results on the LOAD-mutants of ADAM10 (supplementary Figure 2) raise the question whether these intra-species variations of ADAM10 might enhance its function as α-toxin receptor. If LOAD-mutants enhance binding of α-toxin, it would fuel speculations that these variants might promote entry of α-toxin into the CNS through the olfactory route, and neurodegeneration, as recently discussed (von Hoven, Qin et al. 2019).

Binding of α-toxin to cells could involve other molecules, possibly in addition to ADAM10 and lipids. A caveolin-binding motif has been identified in α-toxin (Pany, Vijayvargia et al. 2004). Caveolin appears to be dispensable for α-toxin-dependent cytotoxicity (Husmann, Beckmann et al. 2009), but radiolabeled α-toxin and caveolin co-migrate as a high-molecular weight complex in a two-dimensional separation of cell lysates from toxin-treated HaCaT cells through density gradients and subsequent SDS-PAGE; further, an antibody to multi-ubiquitin precipitated α-toxin-oligomers (von Hoven 2012). Possibly, caveolin and ubiquitination play a role in the traffic of α-toxin.

Advancing knowledge about toxin/receptor interactions may pave the way for development of new therapeutic approaches. Antibodies against α-toxin or decoy liposomes might help fight certain infections by *S. aureus*, including by MRSA (Henry, Neill et al. 2015, Tkaczyk, Kasturirangan et al. 2017). Receptor-targeted drugs could be a valuable alternative. Although GI254023X, a small molecular weight inhibitor of ADAM10 (Hundhausen, Misztela et al. 2003), blocks α-toxin - binding and cytotoxicity *in vitro* (von Hoven, Rivas et al. 2016), its therapeutic use may be obsolete because of potential unwanted effects due to the inhibition of ADAM10’s catalytic activity. The present results might inform the design of peptides or peptido-mimetics (Modell, Blosser et al. 2016, Bruzzoni-Giovanelli, Alezra et al. 2018), to inhibit docking of α–toxin to target cells without interfering with physiological functions of ADAM10.

Animal models do not perfectly reflect human physiology (Drake 2013). Both mouse adapted *S. aureus* strains (Trübe, Hertlein et al. 2019), as well as humanized mouse models, e.g. (Parker 2017, Prince, Wang et al. 2017), may improve *in vivo* studies of *S. aureus* toxins, or infections by this organism. The present results suggest that mutating mouse, rat or guinea pig ADAM10 could enhance the suitability of these animals for studying *in vivo* effects of α-toxin under conditions relevant for humans. It is possible that a combination from the eight residues differing between human (or bovine) and murine ΔPD sequences would yield a larger effect than the single mutation Q666E alone. However, finding all of the 255 possible combinations in a pool of synthetic ADAM10 cDNAs, randomly mutagenized at these eight positions would come down to the *coupon collector’s problem* requiring functional testing of ∼1.500 clones. Yet, because replacement of Q666 in mADAM10 with glutamine significantly increased the efficiency of mADAM10 as α-toxin receptor *in vitro*, it appears quite promising to investigate the effect of this single mutation *in vivo*.

Conversely, it may be desirable to make livestock more resistant to *S. aureus*. Infections with this bacterium are a notorious problem in dairy cows. Reportedly, within-host adaptation includes increased cytotoxicity with increased secretion of α-toxin (Mayer, Kucklick et al. 2021). Therefore, mutation E665Q might enhance resilience of cows to *S. aureus*.

In conclusion, we propose that E665 of bovine ADAM10 is important for the role of this protein as a receptor for α-toxin. The initial interaction of α-toxin monomers with membranes could involve dual binding to both phosphocholine *and* ADAM10. Binding of α-toxin to ADAM10 might involve direct interaction with E665. By inference, the present findings likely hold for hADAM10 and most orthologues. In contrast, mice have evolved a glutamine residue at the corresponding position, rendering mADAM10 a comparably inefficient receptor for α-toxin. This may explain relative tolerance of many murine cells to attack by this pore-forming toxin.

## Materials and Methods

### Antibodies, annexin V, α-toxin and peptides

AB19026 ADAM10 rabbit polyclonal antibody (Immunogen: hADAM10 peptide 732-748) was purchased from Merck; anti-ADAM10 IC1427F was from R&D; anti-HA-Tag 6E2, a mouse mAb, was from CellSignaling (#2367). Annexin V-AlexaFluo647 (A23204) was purchased from Invitrogen. *S. aureus* α-toxin was prepared as described earlier (Palmer, Jürsch et al. 1993). Peptides (Acetyl)-SPELYENIA(Amide) and (Acetyl)SPKLYENIA(Amide), covering wild type or mutant bADAM10 sequence, respectively, were custom-synthesized by ThermoScientific and delivered as lyophilized, trifluoroacetic salts. After slow equilibration to room temperature (RT), we dissolved peptides in a 50% (v/v) aqueous solution of DMSO.

### Plasmid constructs

Expression constructs for the PD or ΔPD of bADAM10 have been described earlier (von Hoven, Rivas et al. 2016), and references therein. Additional truncations, deletions or two compound mutations (changes at position N°2, N°3, N°4 and N°6 or positions N°1, N°5, N°7 and N°8 as shown in Fig. 3B) were custom-synthesized, and provided as inserts in pEX-A258 or pEX- A128 (Eurofins). Plasmids were transformed into DH5α, and plasmid DNA prepared from clones. Inserts were subcloned via XhoI and XbaI sites into pcDNA3 (Invitrogen). Low endotoxin plasmid DNA for transfection experiments was obtained by using kits from Machery&Nagel. For mutation of single nucleotides we employed QuikChange II Site-directed mutagenesis kit (Agilent) or NEBase Changer–Kit (NEB) and primers listed in supplementary Table 1. All constructs were verified by sequencing.

### Cells and culture conditions

The murine keratinocyte cell line PDV and the human keratinocyte cell line HaCaT were cultured as described (von Hoven, Neukirch et al. 2015). HAP1 (near-haploid human cell line, derived from KBM-7 cells (Carette, Raaben et al. 2011) were grown in Iscove’s modified Dulbecco’s medium (IMDM) in the presence of 10% (v/v) FBS and penicillin/streptomycin. HAP1ADAM10KO cells have a frame shift mutation in the ADAM10 gene and were custom made by Horizon using the clustered regularly interspaced short palindromic repeats (CRISPR)/CRISPR-associated 9 (Cas9) system; parental HAP1 cells served as a control. All media and medium additives were from Gibco/Life Technologies.

### Transient transfection

HAP1ADAM10KO cells were transfected with plasmids by electroporation, using the Amaxa system (Lonza) according to the manufacturer’s protocol and nucleofector kit V, program T020).

### ATP-assay

Luminometric measurements of cellular ATP levels were performed as described elsewhere (Haugwitz, Bobkiewicz et al. 2006). In brief, cells were seeded at a density of 2×10^4^ per well of a 96-well tissue culture plate. After 24 h treatments were performed and ATP was measured using the ATP Bioluminescence Assay Kit CLSII from Roche. This assay exploits the fact that luciferase requires ATP to produce light, which was measured with a Lumat 9705 instrument from Berthold.

### Western-blot

For Western blots, cell lysates were mixed with an equal volume of 2 × SDS loading buffer (10% (v/v) glycerol, 5% (v/v) 2-mercaptoethanol, 2% (w/v) SDS and Bromphenol Blue) and heated for 5 min at 95°C. Proteins were separated by SDS/PAGE (10% gel) and transferred onto a nitrocellulose membrane. Subsequently, the membrane was blocked for 1h at RT in 5% (w/v) BSA, or 5% (w/v) dried non-fat skimmed milk powder in TBST (50 μM Tris/HCl, 0.15 M NaCl and 0.1% Tween 20), was followed by incubating the membrane with primary antibody in 5% (w/v) bovine serum albumin (BSA), or 5% (w/v) dried non-fat skimmed milk powder in TBST (overnight at 4°C), washed three times in TBST and incubated with horseradish peroxidase-conjugated second antibody from Cell Signaling Technology (dilution 1:2000) for 1 h at RT. After three washing steps, detection of bound conjugate was achieved by the use of an enhanced chemiluminescence assay (ECL) kit (Roche Applied Science). Images were acquired using a BIO-RAD ChemiDoc imaging system.

### Flow cytometry

For the detection of ADAM10 surface expression, HAP1ADAM10KO cells transfected with plasmids expressing ADAM10 variants or mock transfected cells were detached and counted in a Neubauer chamber. To wash cells, 1×10^5^ cells per sample were spun down at 300 x g for 5 min at 4°C, re-suspended in ice-cold phosphate buffered saline (PBS), spun down again and the supernatant was removed before cells were processed. The pellet was re-suspended in 3% BSA/PBS for 30 min at 4°C (blocking step). After two washing steps as described above, cells were incubated with a fluorescein-isothiocyanate (FITC)-conjugated monoclonal antibody directed against hADAM10 (IC1427F) for 45 min at 4°C protected from light. Subsequently, after two washing steps cells were re-suspended in 500 μl PBS containing 10% FCS and 1% sodium azide. Samples were analyzed using a BD FACS CANTO II and FACS Diva software, as well as FlowJo^TM^.

For analysis of annexin V-binding and propidium iodine (PI) influx, cells were treated or not with α-toxin as indicated in Figure 1A, washed twice with PBS, detached with trypsin, re-suspended in complete culture medium, and washed twice with ice cold PBS. The pellet was re-suspended in 100 μl Annexin-binding-buffer (10 mM HEPES, 140 mM NaCl, 2,5 mM CaCl_2_ pH 7.4), with 1 μl PI stock solution (100 μg/ml) plus 3.5 μl AnnexinV-Alexa647. After incubation for 15 min at RT 400 μl of Annexin-binding buffer were added, and samples were kept on ice before analysis by FACS (see above).

### Binding assay with radioactive α-toxin

To detect α-toxin monomers and oligomers associated with cells we employed a procedure based on the use of internally [^35^S]methionine-radiolabeled α-toxin (K21C), immune-precipitation, SDS-PAGE and fluorography, as published earlier (Husmann, Beckmann et al. 2009). In brief, cells were incubated with 1 μg internally labeled α-toxin for 2 h at 37°C, washed and lysed. Equal amounts of solubilized protein were subjected to immunoprecipitation with polyclonal rabbit antibody to α-toxin, (Sigma). Finally, α-toxin monomers and SDS-stable oligomers were separated by electrophoresis, and detected by fluorography.

### Hemolysis assay

First, a suspension containing 5% RRBC in PBS was prepared from the stock solution containing 50% RRBC in Alsever’s solution (Preclinics). RRBC were incubated with different doses of *S. aureus* α-toxin in the presence or absence of 10 μM ADAM10-peptides for 30 min at 37°C, and hemolysis was assessed by measuring absorption at 415 nm using a BioRad iMark Microplate Reader.

### *In silico* docking of α-toxin to ADAM10

For *in silico* docking of α-toxin to ADAM10 we employed HDOCK (Yan, Tao et al. 2020), and ClusPro (Kozakov, Hall et al. 2017) servers. As receptor, we used the model of the ADAM10 ectodomain including the stalk region shown in Figure 5C. It was created by using the I-TASSER server (Yang, Yan et al. 2015), from a sequence covering both the part of the hADAM10 ectodomain, the structure of which (RCDS PDB 6BDZ), has been published (Seegar, Killingsworth et al. 2017), plus the sequence of the stalk region (see Figure 5A). As ligand, we used the A-chain of 4IDJ (RCSD PDB), a structure of the α-toxin monomer (Foletti, Strop et al. 2013). Based on the assumption that E665 of ADAM10, if directly involved in the binding of α-toxin, would contact the membrane-proximal region of the α-toxin monomer, we selected a model created by HDOCK without restraints, where E665 of ADAM10 was in proximity to S203 of α-toxin. In ClusPro, according restraint settings (i.e. E665 in a distance of 1-5 Å to S203 of α-toxin) led to a model underlying Figure 6; for visualization of the model as shown, we used PyMOL release from 2006, (DeLano, W.); current versions: http://www.pyol.org/pymol.

### Phylogenetic tree of ADAM10

Multiple sequence alignment of ADAM10 protein sequences of different species was performed with Clustal Omega EMBL-EBI (Madeira, Park et al. 2019) using protein sequences in FASTA format (supplementary information). In order to create a phylogenic tree, the alignment file was downloaded in Pearson/FASTA format and uploaded to the online tool iTOL v5 Interactive Tree Of Life (https://itol.embl.de) (Letunic and Bork 2021), rooted to *Danio rerio*, and annotated. The names of species with glutamine, Q, in positions corresponding to residue 666 of mADAM10 are in magenta.

### Statistics

Numerical data displayed in bar graphs represent mean values ± SEM from n ≥ 3 independent experiments. GraphPadPrism8 was employed for selection and application of tests for significance; details are mentioned in the figure legends. Significance was generally assumed at *P* <0.05.

## Acknowledgements

We gratefully acknowledge technical assistance by S. Möckel and S. Nick from the Flow Cytometry Core Facility at IMB Mainz, Germany, and J. Altmeier from FACSlab, Institute for Toxicology, University Medical Center Mainz. We like to thank R. Postina, Johannes Gutenberg-University Mainz, for sharing plasmids and K. Husmann for calculations. Parts of the work have been done in partial fulfillment of the doctoral thesis of DS. The University Medical Center of the Johannes Gutenberg-University Mainz supported this work by a grant to GvH.

## Author contributions

MH and GvH conceived of the study. MM produced ADAM10 constructs. CN, MM and DS performed experiments, acquired and analyzed data. GvH designed experiments, analyzed data, interpreted data and wrote the manuscript. MH analyzed and interpreted data and wrote the manuscript. All authors contributed to manuscript revision, read and approved the submitted version.

## Abbreviations

ADAM10,: A Disintegrin And Metalloprotease;
bADAM10,: bovine ADAM10;
FACS,: fluorescence activated cell sorting;
FITC,: fluorescein-isothiocyanate;
HA,: Hemagglutinin;
hADAM10,: human ADAM10;
HAP1ADAM10KO cells,: haploid human cell line with knock out of ADAM10 by CRISPR/CAS9;
I-TASSER,: Iterative Threading ASSEmbly Refinement;
LOAD,: Late Onset Alzheimer’s Disease;
mADAM10,: murine ADAM10;
PD,: prodomain of ADAM10;
ΔPD,: ADAM10 devoid of the PD;
PI: propidium iodide;
RRBC,: rabbit red blood cells

## References

Anders, A., S. Gilbert, W. Garten, R. Postina and F. Fahrenholz (2001). “Regulation of the alpha-secretase ADAM10 by its prodomain and proprotein convertases.” FASEB J 15(10): 1837–1839. doi:10.1096/fj.01-0007fje.

Berube, B. J. and J. Bubeck Wardenburg (2013). “Staphylococcus aureus alpha-toxin: nearly a century of intrigue.” Toxins (Basel) 5(6): 1140–1166. https://www.ncbi.nlm.nih.gov/pubmed/23888516 https://www.ncbi.nlm.nih.gov/pmc/articles/PMC3717774/pdf/toxins-05-01140.pdf

Bhakdi, S., M. Muhly, S. Korom and F. Hugo (1989). “Release of interleukin-1 beta associated with potent cytocidal action of staphylococcal alpha-toxin on human monocytes.” Infect Immun 57(11): 3512–3519. doi:10.1128/iai.57.11.3512-3519.1989.

Bleibaum, F., A. Sommer, M. Veit, B. Rabe, J. Andra, K. Kunzelmann, C. Nehls, W. Correa, T. Gutsmann, J. Grotzinger, S. Bhakdi and K. Reiss (2019). “ADAM10 sheddase activation is controlled by cell membrane asymmetry.” J Mol Cell Biol 11(11): 979–993. doi:10.1093/jmcb/mjz008.

Bruzzoni-Giovanelli, H., V. Alezra, N. Wolff, C. Z. Dong, P. Tuffery and A. Rebollo (2018). “Interfering peptides targeting protein-protein interactions: the next generation of drugs?” Drug Discovery Today 23(2): 272–285. doi:10.1016/j.drudis.2017.10.016.

Carette, J. E., M. Raaben, A. C. Wong, A. S. Herbert, G. Obernosterer, N. Mulherkar, A. I. Kuehne, P. J. Kranzusch, A. M. Griffin, G. Ruthel, P. Dal Cin, J. M. Dye, S. P. Whelan, K. Chandran and T. R. Brummelkamp (2011). “Ebola virus entry requires the cholesterol transporter Niemann-Pick C1.” Nature 477(7364): 340–343. doi:10.1038/nature10348.

Cassidy, P. S. and S. Harshman (1973). “The binding of staphylococcal 125I-alpha-toxin (B) to erythrocytes.” J Biol Chem 248(15): 5545–5546. https://www.ncbi.nlm.nih.gov/pubmed/4806341

Dragneva, Y., C. D. Anuradha, A. Valeva, A. Hoffmann, S. Bhakdi and M. Husmann (2001). “Subcytocidal attack by staphylococcal alpha-toxin activates NF-kappaB and induces interleukin-8 production.” Infect Immun 69(4): 2630–2635. doi:10.1128/IAI.69.4.2630-2635.2001.

Drake, A. C. (2013). “Of mice and men: what rodent models don’t tell us.” Cell Mol Immunol 10(4): 284–285. doi:10.1038/cmi.2013.21.

Ezekwe, E. A., Jr., C. Weng and J. A. Duncan (2016). “ADAM10 Cell Surface Expression but Not Activity Is Critical for Staphylococcus aureus alpha-Hemolysin-Mediated Activation of the NLRP3 Inflammasome in Human Monocytes.” Toxins (Basel) 8(4): 95. doi:10.3390/toxins8040095.

Foletti, D., P. Strop, L. Shaughnessy, A. Hasa-Moreno, M. G. Casas, M. Russell, C. Bee, S. Wu, A. Pham, Z. Zeng, J. Pons, A. Rajpal and D. Shelton (2013). “Mechanism of action and in vivo efficacy of a human-derived antibody against Staphylococcus aureus alpha-hemolysin.” J Mol Biol 425(10): 1641–1654. doi:10.1016/j.jmb.2013.02.008.

Galdiero, S. and E. Gouaux (2004). “High resolution crystallographic studies of alpha-hemolysin-phospholipid complexes define heptamer-lipid head group interactions: implication for understanding protein-lipid interactions.” Protein Sci 13(6): 1503–1511. doi:10.1110/ps.03561104.

Haugwitz, U., W. Bobkiewicz, S. R. Han, E. Beckmann, G. Veerachato, S. Shaid, S. Biehl, K. Dersch, S. Bhakdi and M. Husmann (2006). “Pore-forming Staphylococcus aureus alpha-toxin triggers epidermal growth factor receptor-dependent proliferation.” Cell Microbiol 8(10): 1591–1600. doi:10.1111/j.1462-5822.2006.00733.x.

Henry, B. D., D. R. Neill, K. A. Becker, S. Gore, L. Bricio-Moreno, R. Ziobro, M. J. Edwards, K. Muhlemann, J. Steinmann, B. Kleuser, L. Japtok, M. Luginbuhl, H. Wolfmeier, A. Scherag, E. Gulbins, A. Kadioglu, A. Draeger and E. B. Babiychuk (2015). “Engineered liposomes sequester bacterial exotoxins and protect from severe invasive infections in mice.” Nat Biotechnol 33(1): 81–88. doi:10.1038/nbt.3037.

Hildebrand, A., M. Pohl and S. Bhakdi (1991). “Staphylococcus aureus alpha-toxin. Dual mechanism of binding to target cells.” J Biol Chem 266(26): 17195–17200. https://www.ncbi.nlm.nih.gov/pubmed/1894613

Hundhausen, C., D. Misztela, T. A. Berkhout, N. Broadway, P. Saftig, K. Reiss, D. Hartmann, F. Fahrenholz, R. Postina, V. Matthews, K. J. Kallen, S. Rose-John and A. Ludwig (2003). “The disintegrin-like metalloproteinase ADAM10 is involved in constitutive cleavage of CX3CL1 (fractalkine) and regulates CX3CL1-mediated cell-cell adhesion.” Blood 102(4): 1186–1195. doi:10.1182/blood-2002-12-3775.

Husmann, M., E. Beckmann, K. Boller, N. Kloft, S. Tenzer, W. Bobkiewicz, C. Neukirch, H. Bayley and S. Bhakdi (2009). “Elimination of a bacterial pore-forming toxin by sequential endocytosis and exocytosis.” FEBS Lett 583(2): 337–344. doi:10.1016/j.febslet.2008.12.028.

Inoshima, I., N. Inoshima, G. A. Wilke, M. E. Powers, K. M. Frank, Y. Wang and J. Bubeck Wardenburg (2011). “A Staphylococcus aureus pore-forming toxin subverts the activity of ADAM10 to cause lethal infection in mice.” Nat Med 17(10): 1310–1314. doi:10.1038/nm.2451.

Jonas, D., I. Walev, T. Berger, M. Liebetrau, M. Palmer and S. Bhakdi (1994). “Novel path to apoptosis: small transmembrane pores created by staphylococcal alpha-toxin in T lymphocytes evoke internucleosomal DNA degradation.” Infect Immun 62(4): 1304–1312. https://www.ncbi.nlm.nih.gov/pubmed/8132337 https://iai.asm.org/content/iai/62/4/1304.full.pdf

Kim, M., J. Suh, D. Romano, M. H. Truong, K. Mullin, B. Hooli, D. Norton, G. Tesco, K. Elliott, S. L. Wagner, R. D. Moir, K. D. Becker and R. E. Tanzi (2009). “Potential late-onset Alzheimer’s disease-associated mutations in the ADAM10 gene attenuate {alpha}-secretase activity.” Hum Mol Genet 18(20): 3987–3996. doi:10.1093/hmg/ddp323.

Kozakov, D., D. R. Hall, B. Xia, K. A. Porter, D. Padhorny, C. Yueh, D. Beglov and S. Vajda (2017). “The ClusPro web server for protein-protein docking.” Nat Protoc 12(2): 255–278. doi:10.1038/nprot.2016.169.

Letunic, I. and P. Bork (2021). “Interactive Tree Of Life (iTOL) v5: an online tool for phylogenetic tree display and annotation.” Nucleic Acids Research 49(W1): W293–W296. doi:10.1093/nar/gkab301.

Madeira, F., Y. M. Park, J. Lee, N. Buso, T. Gur, N. Madhusoodanan, P. Basutkar, A. R. N. Tivey, S. C. Potter, R. D. Finn and R. Lopez (2019). “The EMBL-EBI search and sequence analysis tools APIs in 2019.” Nucleic Acids Res 47(W1): W636–W641. doi:10.1093/nar/gkz268.

Maurer, K., T. Reyes-Robles, F. Alonzo, 3rd, J. Durbin, V. J. Torres and K. Cadwell (2015). “Autophagy mediates tolerance to Staphylococcus aureus alpha-toxin.” Cell Host Microbe 17(4): 429–440. doi:10.1016/j.chom.2015.03.001.

Mayer, K., M. Kucklick, H. Marbach, M. Ehling-Schulz, S. Engelmann and T. Grunert (2021). “Within-Host Adaptation of Staphylococcus aureus in a Bovine Mastitis Infection Is Associated with Increased Cytotoxicity.” Int J Mol Sci 22(16). doi:10.3390/ijms22168840.

Modell, A. E., S. L. Blosser and P. S. Arora (2016). “Systematic Targeting of Protein-Protein Interactions.” Trends in Pharmacological Sciences 37(8): 702–713. doi:10.1016/j.tips.2016.05.008.

Noskin, G. A., R. J. Rubin, J. J. Schentag, J. Kluytmans, E. C. Hedblom, M. Smulders, E. Lapetina and E. Gemmen (2005). “The burden of Staphylococcus aureus infections on hospitals in the United States: an analysis of the 2000 and 2001 Nationwide Inpatient Sample Database.” Arch Intern Med 165(15): 1756–1761. doi:10.1001/archinte.165.15.1756.

Palmer, M., R. Jürsch, U. Weller, A. Valeva, K. Hilgert, M. Kehoe and S. Bhakdi (1993). “Staphylococcus aureus alpha-toxin. Production of functionally intact, site-specifically modifiable protein by introduction of cysteine at positions 69, 130, and 186.” J Biol Chem 268(16): 11959–11962. https://www.ncbi.nlm.nih.gov/pubmed/8505320

Pany, S., R. Vijayvargia and M. V. Krishnasastry (2004). “Caveolin-1 binding motif of alpha-hemolysin: its role in stability and pore formation.” Biochem Biophys Res Commun 322(1): 29–36. doi:10.1016/j.bbrc.2004.07.073.

Parker, D. (2017). “Humanized Mouse Models of Staphylococcus aureus Infection.” Front Immunol 8: 512. doi:10.3389/fimmu.2017.00512.

Popov, L. M., C. D. Marceau, P. M. Starkl, J. H. Lumb, J. Shah, D. Guerrera, R. L. Cooper, C. Merakou, D. M. Bouley, W. Meng, H. Kiyonari, M. Takeichi, S. J. Galli, F. Bagnoli, S. Citi, J. E. Carette and M. R. Amieva (2015). “The adherens junctions control susceptibility to Staphylococcus aureus alpha-toxin.” Proc Natl Acad Sci U S A 112(46): 14337–14342. doi:10.1073/pnas.1510265112.

Prince, A., H. Wang, K. Kitur and D. Parker (2017). “Humanized Mice Exhibit Increased Susceptibility to Staphylococcus aureus Pneumonia.” J Infect Dis 215(9): 1386–1395. doi:10.1093/infdis/jiw425.

Schwiering, M., A. Brack, R. Stork and N. Hellmann (2013). “Lipid and phase specificity of alpha-toxin from S. aureus.” Biochim Biophys Acta 1828(8): 1962–1972. doi:10.1016/j.bbamem.2013.04.005.

Seegar, T. C. M., L. B. Killingsworth, N. Saha, P. A. Meyer, D. Patra, B. Zimmerman, P. W. Janes, E. Rubinstein, D. B. Nikolov, G. Skiniotis, A. C. Kruse and S. C. Blacklow (2017). “Structural Basis for Regulated Proteolysis by the alpha-Secretase ADAM10.” Cell 171(7): 1638–1648 e1637. doi:10.1016/j.cell.2017.11.014.

Song, L. Z., M. R. Hobaugh, C. Shustak, S. Cheley, H. Bayley and J. E. Gouaux (1996). “Structure of staphylococcal alpha-hemolysin, a heptameric transmembrane pore.” Science 274(5294): 1859–1866. doi:DOI 10.1126/science.274.5294.1859.

Suh, J., S. H. Choi, D. M. Romano, M. A. Gannon, A. N. Lesinski, D. Y. Kim and R. E. Tanzi (2013). “ADAM10 missense mutations potentiate beta-amyloid accumulation by impairing prodomain chaperone function.” Neuron 80(2): 385–401. doi:10.1016/j.neuron.2013.08.035.

Tkaczyk, C., S. Kasturirangan, A. Minola, O. Jones-Nelson, V. Gunter, Y. Y. Shi, K. Rosenthal, V. Aleti, E. Semenova, P. Warrener, D. Tabor, C. K. Stover, D. Corti, G. Rainey and B. R. Sellman (2017). “Multimechanistic Monoclonal Antibodies (MAbs) Targeting Staphylococcus aureus Alpha-Toxin and Clumping Factor A: Activity and Efficacy Comparisons of a MAb Combination and an Engineered Bispecific Antibody Approach.” Antimicrob Agents Chemother 61(8). doi:10.1128/AAC.00629-17.

Trübe, P., T. Hertlein, D. M. Mrochen, D. Schulz, I. Jorde, B. Krause, J. Zeun, S. Fischer, S. A. Wolf, B. Walther, T. Semmler, B. M. Broker, R. G. Ulrich, K. Ohlsen and S. Holtfreter (2019). “Bringing together what belongs together: Optimizing murine infection models by using mouse-adapted Staphylococcus aureus strains.” Int J Med Microbiol 309(1): 26–38. doi:10.1016/j.ijmm.2018.10.007.

Valeva, A., N. Hellmann, I. Walev, D. Strand, M. Plate, F. Boukhallouk, A. Brack, K. Hanada, H. Decker and S. Bhakdi (2006). “Evidence that clustered phosphocholine head groups serve as sites for binding and assembly of an oligomeric protein pore.” J Biol Chem 281(36): 26014–26021. doi:10.1074/jbc.M601960200.

Valeva, A., I. Walev, M. Pinkernell, B. Walker, H. Bayley, M. Palmer and S. Bhakdi (1997). “Transmembrane beta-barrel of staphylococcal alpha-toxin forms in sensitive but not in resistant cells.” Proc Natl Acad Sci U S A 94(21): 11607–11611. https://www.ncbi.nlm.nih.gov/pubmed/9326657 https://www.ncbi.nlm.nih.gov/pmc/articles/PMC23553/pdf/pq011607.pdf

Virreira Winter, S., A. Zychlinsky and B. W. Bardoel (2016). “Genome-wide CRISPR screen reveals novel host factors required for Staphylococcus aureus alpha-hemolysin-mediated toxicity.” Scientific Reports 6. doi:i:ARTN 24242 10.1038/srep24242.

von Hoven, G. (2012). “Induktion von Pro-Autophagie-Signalen durch einen extra-oder intrazellulären Alpha-Toxin-Angriff.” doi:i:10.25358/openscience-4414.

von Hoven, G., C. Neukirch, M. Meyenburg, S. Füser, M. B. Petrivna, A. J. Rivas, A. Ryazanov, R. J. Kaufman, R. V. Aroian and M. Husmann (2015). “eIF2alpha Confers Cellular Tolerance to S. aureus alpha-Toxin.” Front Immunol 6: 383. doi:10.3389/fimmu.2015.00383.

von Hoven, G., Q. Qin, C. Neukirch, M. Husmann and N. Hellmann (2019). “Staphylococcus aureus alpha-toxin: small pore, large consequences.” Biol Chem 400(10): 1261–1276. doi:10.1515/hsz-2018-0472.

von Hoven, G., A. J. Rivas, C. Neukirch, S. Klein, C. Hamm, Q. Qin, M. Meyenburg, S. Füser, P. Saftig, N. Hellmann, R. Postina and M. Husmann (2016). “Dissecting the role of ADAM10 as a mediator of Staphylococcus aureus alpha-toxin action.” Biochem J 473(13): 1929–1940. doi:10.1042/BCJ20160062.

Walev, I., E. Martin, D. Jonas, M. Mohamadzadeh, W. Müller-Klieser, L. Kunz and S. Bhakdi (1993). “Staphylococcal alpha-toxin kills human keratinocytes by permeabilizing the plasma membrane for monovalent ions.” Infect Immun 61(12): 4972–4979. https://www.ncbi.nlm.nih.gov/pubmed/8225571 https://iai.asm.org/content/iai/61/12/4972.full.pdf

Walev, I., M. Palmer, E. Martin, D. Jonas, U. Weller, H. Höhn-Bentz, M. Husmann and S. Bhakdi (1994). “Recovery of human fibroblasts from attack by the pore-forming alpha-toxin of Staphylococcus aureus.” Microb Pathog 17(3): 187–201. doi:10.1006/mpat.1994.1065.

Wilke, G. A. and J. Bubeck Wardenburg (2010). “Role of a disintegrin and metalloprotease 10 in Staphylococcus aureus alpha-hemolysin-mediated cellular injury.” Proc Natl Acad Sci U S A 107(30): 13473–13478. doi:10.1073/pnas.1001815107.

Yan, Y., H. Tao, J. He and S. Y. Huang (2020). “The HDOCK server for integrated protein-protein docking.” Nat Protoc 15(5): 1829–1852. doi:10.1038/s41596-020-0312-x.

Yang, J. Y., R. X. Yan, A. Roy, D. Xu, J. Poisson and Y. Zhang (2015). “The I-TASSER Suite: protein structure and function prediction.” Nature Methods 12(1): 7–8. doi:10.1038/nmeth.3213.

Ziesemer, S., N. Möller, A. Nitsch, C. Müller, A. G. Beule and J. P. Hildebrandt (2019). “Sphingomyelin Depletion from Plasma Membranes of Human Airway Epithelial Cells Completely Abrogates the Deleterious Actions of S. aureus Alpha-Toxin.” Toxins (Basel) 11(2). doi:10.3390/toxins11020126.

