## Supplemental material for "Glutamine 666 renders murine ADAM10 an inefficient *S. aureus* α-toxin receptor"

Supplementary Figure 1

```
        .........10 .........20 .........30 .........40 .........50
hADAM10 MVLLRVLILL LSWAAGMGGQ YGNPLNKYIR HYEGLSYNVD SLHQKHQRAK
bADAM10 MVLLRVLILL LSWVAGLGGQ YGNPLNKYIR HYEGLSYDVD SLHQKHQRAK
rADAM10 MILIKTVVLI FI------GQ YGNPLNKYIR HYEGLSYSVD SLHQKHQRAK
mADAM10 MVLPTVLILL LSWAAGLGGQ YGNPLNKYIR HYEGLSYNVD SLHQKHQRAK

        .........60 .........70 .........80 .........90 .........100
hADAM10 RAVSHEDQFL RLDFHAHGRH FNLRMKRDTS LFSDEFKVET SNKVLDYDTS
bADAM10 RAVSHEDQFL RLDFHAHGRH FNLRMKRDTS LFSEEFRVET SNAVLDYDTS
rADAM10 RAVSHEDQFL RLDFHAHGRH FNLRMKRDTS LFSDDFKVET SNKVLDYDTS
mADAM10 RAVSHEDQFL LLDFHAHGRQ FNLRMKRDTS LFSDEFKVET SNKVLDYDTS

        .........110 .........120 .........130 .........140 .........150
hADAM10 HIYTGHIYGE EGSFSHGSVI DGRFEGFIQT RGGTFYVEPA ERYIKDRTLP
bADAM10 HIYTGHIYGE EGSFSHGSVI DGRFEGFIQT HGGTFYVEPA ERYIKDRTLP
rADAM10 HIYTGHIYGE EGSFSHGSVI DGRFEGFIQT RGGTFYIEPA ERYIKDRTLP
mADAM10 HIYTGHIYGE EGSFSHGSVI DGRFEGFIKT RGGTFYIEPA ERYIKDRILP

        .........160 .........170 .........180 .........190 .........200
hADAM10 FHSVIYHEDD INYPHKYGPQ GGCADHSVFE RMRKYQMTGV EEVTQIPQEE
bADAM10 FHSVIYHEDD IKYPHKYGPQ GGCADHSVFE RMRKYQMTGV EEVTQTPQEK
rADAM10 FHSVIYHEDD INYPHKYGPQ GGCADHSVFE RMKKYQMTGV EEVTQTPQEE
mADAM10 FHSVIYHEDD INYPHKYGPQ GGCADHSVFE RMRKYQMTGV EEGARAHPEK

        .........210 .........220 .........230 .........240 .........250
hADAM10 HAA-NGPELL RKKRTTSAEK NTCQLYIQTD HLFFKYYGTR EAVIAQISSH
bADAM10 HAI-NGPELL RKKRTTVAEK NTCQLYIQTD HLFFKYYGTR EAVIAQISSH
rADAM10 HAA-NGPELL RKKRTTSAEK NTCQLYIQTD HLFFKYYGTR EAVIAQISSH
mADAM10 HAASSGPELL RKKRTTLAER NTCQLYIQTD HLFFKYYGTR EAVIAQISSH

        .........260 .........270 .........280 .........290 .........300
hADAM10 VKAIDTIYQT TDFSGIRNIS FMVKRIRINT TADEKDPTNP FRFPNIGVEK
bADAM10 VKAIDTIYQT TDFSGIRNIS FMVKRIRINT TADEKDPTNP FRFPNIGVEK
rADAM10 VKAIDTIYQT TDFSGIRNIS FMVKRIRINT TADEKDPTNP FRFPNIGVEK
mADAM10 VKAIDTIYQT TDFSGIRNIS FMVKRIRINT TSDEKDPTNP FRFPNIGVEK

        .........310 .........320 .........330 .........340 .........350
hADAM10 FLELNSEQNH DDYCLAYVFT DRDFDDGVLG LAWVGAPSGS SGGICEKSKL
bADAM10 FLELNSEQNH DDYCLAYVFT DRDFDDGVLG LAWVGAPSGS SGGICEKSKL
rADAM10 FLELNSEQNH DDYCLAYVFT DRDFDDGVLG LAWVGAPSGS SGGICEKSKL
mADAM10 FLELNSEQNH DDYCLAYVFT DRDFDDGVLG LAWVGAPSGS SGGICEKSKL

        .........360 .........370 .........380 .........390 .........400
hADAM10 YSDGKKKSLN TGIITVQNYG SHVPPKVSHI TFAHEVGHNF GSPHDSGTEC
bADAM10 YSDGKKKSLN TGIITVQNYG SHVPPKVSHI TFAHEVGHNF GSPHDSGTEC
rADAM10 YSDGKKKSLN TGIITVQNYG SHVPPKVSHI TFAHEVGHNF GSPHDSGTEC
mADAM10 YSDGKKKSLN TGIITVQNYG SHVPPKVSHI TFAHEVGHNF GSPHDSGTEC

        .........410 .........420 .........430 .........440 .........450
hADAM10 TPGESKNLGQ KENGNYIMYA RATSGDKLNN NKFSLCSIRN ISQVLEKKRN
bADAM10 TPGESKNLGQ KENGNYIMYA RATSGDKLNN NKFSLCSIRN ISQVLEKKRN
rADAM10 TPGESKNLGQ KENGNYIMYA RATSGDKLNN NKFSLCSIRN ISQVLEKKRN
mADAM10 TPGESKNLGQ KENGNYIMYA RATSGDKLNN NKFSLCSIRN ISQVLEKKRN

        .........460 .........470 .........480 .........490 .........500
hADAM10 NCFVESGQPI CGNGMVEQGE ECDCGYSDQC KDECCFDANQ PEGRKCKLKP
bADAM10 NCFVESGQPI CGNGMVEQGE ECDCGYSDQC KDECCYDANQ PEGKKCKLKP
rADAM10 NCFVESGQPI CGNGMVEQGE ECDCGYSDQC KDECCFDANQ PEGKKCKLKP
mADAM10 NCFVESGQPI CGNGMVEQGE ECDCGYSDQC KDDCCFDANQ PEGKKCKLKP
```

```
         .........510.........520.........530.........540.........550
hADAM10  GKQCSPSQGP CCTAQCAFKS KSEKCRDDSD CAREGICNGF  TALCPASDPK
bADAM10  GKQCSPSQGP CCTAHCAFKS KTEKCRDDSD CAKEGICNGI  TALCPASDPK
rADAM10  GKQCSPSQGP CCTAQCAFKS KSEKCRDDSD CAREGICNGF  TALCPASDPK
mADAM10  GKQCSPSQGP CCTAQCAFKS KSEKCRDDSD CAKEGICNGF  TALCPASDPK

         .........560.........570.........580.........590.........600
hADAM10  PNFTDCNRHT QVCINGQCAG SICEKYGLEE CTCASSDGKD DKELCHVCCM
bADAM10  PNFTDCNRHT QVCINGQCAG SICEKHGLEE CTCASSDGKD DKELCHVCCM
rADAM10  PNFTDCNRQT QVCINGQCAG SICEKYGLEE CTCASSDGKD DKELCHVCCM
mADAM10  PNFTDCNRHT QVCINGQCAG SICEKYDLEE CTCASSDGKD DKELCHVCCM

         .........610.........620.........630.........640.........650
hADAM10  KKMDPSTCAS TGSVQWSRHF SGRTITLQPG SPCNDFRGYC DVFMRCRLVD
bADAM10  KKMEPSTCAS TGSVQWNKYF LGRTITLQPG SPCNDFRGYC DVFMRCRLVD
rADAM10  KKMEPSTCAS TGSVQWSKQF SGQTITLQPG SPCNDFRGYC DVFMRCRLVD
mADAM10  KKMAPSTCAS TGSLQWSKQF SGRTITLQPG SPCNDFRGYC DVFMRCRLVD

                              ⬇
         .........660.........670.........680.........690.........700
hADAM10  ADGPLARLKK AIFSPELYEN IAEWIVAHWW AVLLMGIALI MLMAGFIKIC
bADAM10  ADGPLARLKK AIFSPELYEN IAEWIVAYWW AVLLMGIALI MLMAGFIKIC
rADAM10  ADGPLARLKK AIFSPELYEN IAEWIVAHWW AVLLMGIALI MLMAGFIKIC
mADAM10  ADGPLARLKK AIFSPQLYEN IAEWIVAHWW AVLLMGIALI MLMAGFIKIC

         .........710.........720.........730.........740.........
hADAM10  SVHTPSSNPK LPPPKPLPGT LKRRRPPQPI QQPQRQRPRE SYQMGHMRR
bADAM10  SVHTPSSNPK LPPPKPLPGT LKRRRPPQPI QQPQRQRPRE SYQMGHMRR
rADAM10  SVHTPSSNPK LPPPKPLPGT LKRRRPPQPI QQPQRQRPRE SYQMGHMRR
mADAM10  SVHTPSSNPK LPPPKPLPGT LKRRRPPQPI QQPPRQRPRE SYQMGHMRR
```

Unconserved 0 1 2 3 4 5 6 7 8 9 10 Conserved

**Suppl. Fig. 1: Alignment of ADAM10 amino acid sequences (human, bovine, rabbit and mouse).** Alignment of sequences was achieved with PRALINE, a multiple sequence alignment toolbox (Simossis, VA and Heringa J, Nucleic Acids Research, 2005; https://doi.org/10.1093/nar/gki390). For sources of sequences, see supplementary information below. Note that there is a shift in numbering of residues due to the additional residue (serine) following residue 203 in the mouse sequence; beyond this point, numbers indicate residues of the mouse sequence. Overall conservation is very high (see color code), but there is a stretch of six residues (193-198) in the prodomain (PD), differing between the murine sequence as compared to rabbit, bovine or human sequence, which however proved to be irrelevant for different activity of mADAM10 as α-toxin-receptor. Instead, a single, rather conservative change (E665 of bADAM10 to Q666 in mADAM10; blue arrow) proved to be critical (see main text). The structure of the ADAM10 ectodomain including the stalk region, which is depicted in Fig. 5C of the manuscript, and was used for *in silico* docking to α-toxin, includes residues 220 through 674.

Supplementary Figure 2

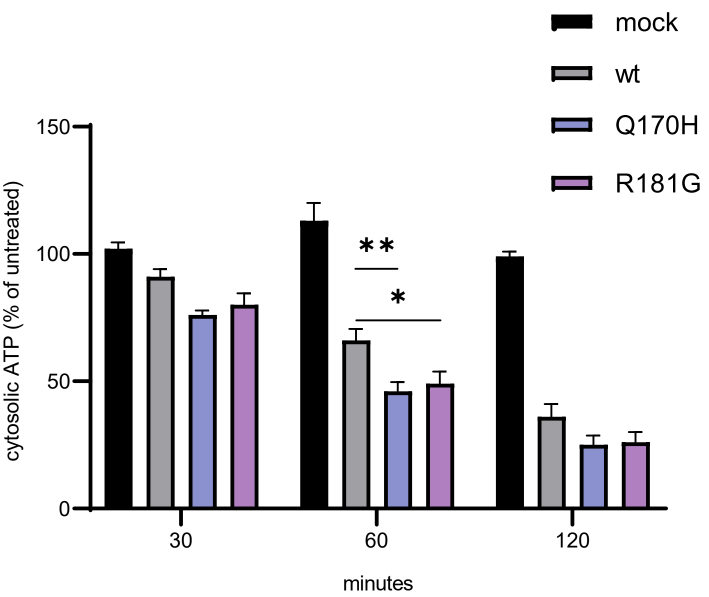


**Suppl. Fig. 2: LOAD-mutants of ADAM10 might confer increased susceptibility to α-toxin**

HAP1ADAM10KO-cells were transiently transfected with expression plasmids for wild type bADAM10, or LOAD mutants (created in the bADAM10 background), as indicated in the figure; mock-transfections were without plasmids. Transfected cell populations were treated, or not, with α-toxin (500 ng/ml) and incubated under culture conditions for the times indicated. Cell lysates were analysed for ATP content by a luminometric assay (see main text). Data are relative ATP levels in toxin-treated vs. untreated cells and represent mean values from ≥ four independent experiments. Statistical significance was assessed with a two-way ANOVA and Tukey´s multiple comparison test. * indicates *P*-value of 0.029, ** of 0.007.

Supplementary Figure 3

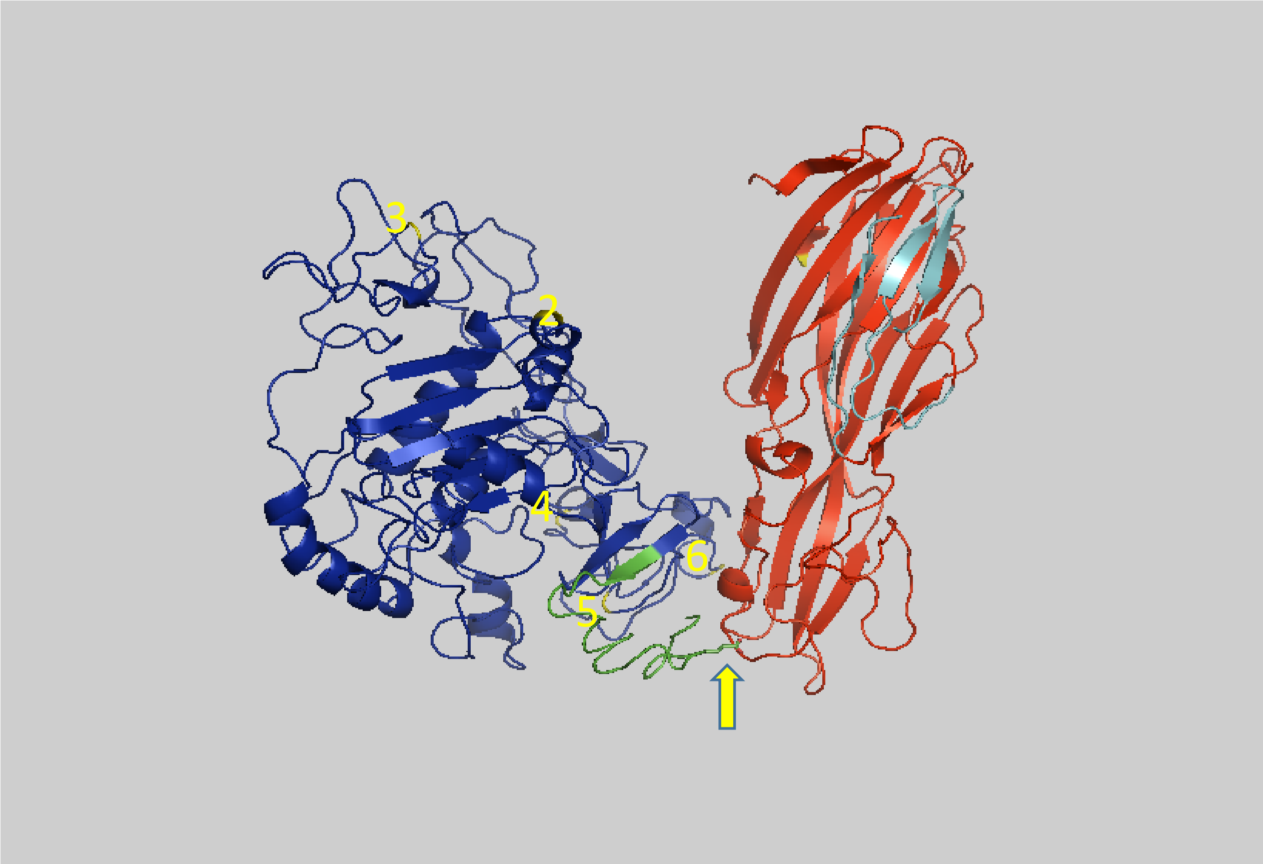


**Suppl. Fig. 3:** The location of some mutations of ADAM10, which we analysed in this work, are indicated by yellow numbers (2, 3, 4, 5 and 6; for details see Fig.3B). The ADAM10 ectodomain is in blue with the stalk region in green, the α-toxin monomer is in red with the pre-stem region highlighted in cyan. None of these mutations affected the function of ADAM10 as a mediator of α-toxin dependent cytotoxicity; and according to this hypothetical model, none of them (including N°6, which is located in a part of the molecule pointing away from the plane of the image) is in proximity to α-toxin. In contrast, residue E665 in the ADAM10 stalk region (yellow arrow) could contribute to direct interaction. Mutations N°1 and N°8 are not shown, because the residues are not part of the structure.

Supplementary Table 1

Primer for mutagenesis

| ADAM10-variant | Primer A 5'-3' | Primer B 5'-3' | Kit |
|---|---|---|---|
| bADAM10 N°8 Q734P | GGGCCTCTGACGCGGGGGCTGTTGAAT | ATTCAACAGCCCCCGCGTCAGAGGCCC | Agilent |
| bADAM10 N°7 E665Q | CAGCTATGTTTTCATAGAGCTGTGGACTGAAAATTGCTTTTTTAAG | CTTAAAAAAGCAATTTTCAGTCCACAGCTCTATGAAAACATAGCTG | Agilent |
| bADAM10 N°7 E665D | TTTCAGTCCAGACCTCTATGAAAAC | ATTGCTTTTTTAAGCCTC | NEB |
| bADAM10 N°7 E665K | TTTCAGTCCAAAGCTCTATGAAAAC | ATTGCTTTTTTAAGCCTCG | NEB |
| bADAM10 NLNN | AACAACGCAATTTTCAGTCCAGAGC | AAGGTTCGCTAGAGGACCATCAGC | NEB |
| bADAM10 NLNNQ | TTTCAGTCCACAGCTCTATGAAAAC | ATTGCGTTGTTAAGGTTC | NEB |

Supplementary Information

ADAM10 Sequences for phylogenetic analysis of ADAM10*

>NP_001152786.1 [Danio rerio], >NP_001083912.1 [Xenopus laevis], >NP_776921.1, >XP_039348190.1 [Mauremys reevesii], >XP_ 040456530.1 [Falco naumanni], >XP_036698615.1 [Balaenotera musculus], [**Bos taurus**], >AAC51766.1 [**Homo sapiens**], >XP_027781569.1 [Marmota flaviventris], >XP_005316677.1 [Ictidomys tridecemlineatus], >XP_002718256.1 [**Oryctolagus cuniculus**], >XP_003462109.2 [Cavia porcellus], >XP_033612141.1 [Fukomys damarensis], >XP_021092820.1 [Heterocephalus glaber], >JAV38388.1 [Castor canadensis], >XP_005075524.1 [Mesocricetus auratus], >NP_062127.1 [Rattus norvegicus] and >NP_031425.2 [**Mus musculus**].

*Sequences in bold face were used for alignment in supplementary Figure 1.
